# Alpha-B-Crystallin overexpression is sufficient to promote tumorigenesis and metastasis in mice

**DOI:** 10.1101/2022.07.27.501790

**Authors:** Behnam Rashidieh, Amanda Louise Bain, Simon Manuel Tria, Sowmya Sharma, Cameron Allan Stewart, Jacinta Ley Simmons, Pirjo M. Apaja, Pascal H.G. Duijf, John Finnie, Kum Kum Khanna

## Abstract

αB-Crystallin is a heat shock chaperone protein which binds to misfolded proteins to prevent their aggregation. It is overexpressed in a wide-variety of cancers. Previous studies using human cancer cell lines and human xenograft models have reported tumor suppressor or tumor promoter (oncogene) roles for αB-Crystallin depending on cellular context and environmental conditions. To determine the causal relationship between *CRYAB* overexpression and cancer, we generated a *Cryab* overexpression knock-in mouse model. This model revealed that constitutive overexpression of *Cryab* results in the formation of a variety of lethal spontaneous primary and metastatic tumors in mice. *In vivo*, the overexpression of *Cryab* correlated with the upregulation of epithelial-to-mesenchymal (EMT) markers, angiogenesis and some oncogenic proteins including Basigin. *In vitro*, using E1A/Ras transformed mouse embryonic fibroblasts (MEFs), we observed that the overexpression of *Cryab* led to the promotion of cell survival via upregulation of Akt signaling and downregulation of pro-apoptotic pathway mediator JNK, with subsequent attenuation of apoptosis as assessed by cleaved caspase-3. Overall, through the generation and characterization of *Cryab* overexpression model, we provide evidence supporting the role of αB-Crystallin as an oncogene, where its upregulation is sufficient to induce tumors, promote cell survival and inhibit apoptosis.

## Introduction

Cancer is a complex genetic disease that stems from the mutations of various genes. These mutations lead to the hallmarks of cancer which favor survival, angiogenesis, invasion, and metastasis^1^. However, the mechanisms that create these advantages also lead to challenges such as endoplasmic reticulum (ER) stress, oxidative stress and hypoxia that can stimulate cell death^2,3^. Cancer cells must overcome these stress responses to develop from a benign tumor into invasive metastatic cancer. Cancer cells proliferate rapidly and synthesize numerous proteins which accumulate and aggregate in the ER leading to ER stress. In normal cells, prolonged ER stress consequently leads to apoptosis and therefore, cancer cells need to overcome this threat^4^. To reduce ER stress, several tumors can upregulate the unfolded protein response (UPR) pathway that helps to preserve intracellular protein homeostasis by increasing the expression of protein folding chaperones including the αB-Crystallin protein encoded by the *CRYAB* gene^2^. The UPR upregulates αB-Crystallin in response to various stimuli such as heat-shock and oxidative stress^2, 5^. Through oligomerizing with other heat-shock proteins, αB-Crystallin allows misfolded or unfolded proteins to be sequestered and prevents their detrimental aggregation which would otherwise create a harmful environment.

The expression of *CRYAB* has been thoroughly investigated in the context of a wide range of cancers where it has been validated as a prognostic marker^6^. In clear cell and papillary type renal cell carcinoma and colorectal cancer, *CRYAB* is used as a marker for higher tumor stage and distant metastases, and in osteosarcoma, it is a marker for increased metastases and poor chemotherapy response^6^. In breast cancer, *CRYAB* is a marker for aggressive behavior in triple-negative basal-like breast cancer and mammary metaplastic carcinoma and its overexpression is associated with the presence of lymph nodal and brain metastasis and relapse^7^. *CRYAB* is also categorized as a marker for lower overall survival in cancers such as renal cell carcinoma, osteosarcoma, colorectal, hepatocellular carcinoma, gastric, ovarian, and non-small cell lung cancer^6^.

At the molecular level, *CRYAB* has been shown to disrupt apoptosis through both the intrinsic and extrinsic pathways^2^ via inhibition of pro-apoptotic Bcl-2 family proteins including Bax and Bcl-xs which contribute to caspase 3 activation. In human mammary epithelial cells, the overexpression of *CRYAB* led to disruption of normal mammary acinar morphology and induction of neoplastic changes^6^. These phenotypes resulted from the activation of survival pathways such as p38, AKT and ERK with *CRYAB* overexpression which could in part be rescued through inhibition of the MEK/ERK pathway^6^. Furthermore, breast cancer cells can induce VEGF expression in co-cultured endothelial cells via activation of UPR and its downstream effector *CRYAB* which protects VEGF from proteolytic degradation^8^.

Despite multiple studies demonstrating a role for *CRYAB* in promoting tumorigenesis *in vitro*, it remains undetermined whether there is a causal link between *CRYAB* overexpression and cancer formation. Here, we demonstrate for the first time, using a transgenic mouse model, that overexpression of *Cryab* is sufficient for *de novo* tumorigenesis. We report that the overexpression of *Cryab* causes high incidence (near 50%) of spontaneous tumor formation with a wide-spectrum of highly proliferative primary and metastatic tumors. Notably, *Cryab* overexpression significantly increased tumor load in a carcinogen-induced tumor model. Using the mouse embryonic fibroblasts (MEFs) from this mouse model, we show that *Cryab* overexpression alters multiple signaling pathways, particularly those related to apoptosis, survival, and metastasis which have potential implications for tumor initiation and therapy development.

## Results

### *CRYAB* is frequently gained and overexpression is associated with poor patient survival in multiple types of cancers

To identify the pattern of *CRYAB* expression levels and copy number alterations in cancer, we performed pan-cancer analyses using TCGA datasets. A comparison of SNP array data revealed that *CRYAB* is gained or amplified in many cancers such as B-Cell neoplasms, non-small cell lung cancer, glioblastoma, melanoma, colorectal, bladder, pancreatic, breast, endometrial, renal, thyroid cancer, and sarcoma (Fig S1a). Additionally, comprehensive survival analyses showed significant associations between high *CRYAB* expression and poor overall patient survival in the vast majority of cancer types, particularly in gastric, bladder, lung cancers. (Fig S1b).

Next, we investigated links between *CRYAB* expression and different features of cancers such as the dysregulation of signaling pathways, angiogenesis, increased proliferation and stemness, neoantigen production, immunoregulation and, tumor infiltration. Notably, this revealed the possible relationship between *CRYAB* expression and dysregulated signaling in different cancer types. For instance, in breast cancer (n=753) there is a positive correlation between *CRYAB* expression and *Ras/MAPK* (r=0.1863, p=2.6 x 10–07); *PI3K/Akt* (r=0.1904, p=1.4 x 10–07); epithelial–mesenchymal transition (EMT) score (r=0.2750, p=1.6 x 10–14); apoptosis (r=0.1238, p=0.0007); autophagy (r=0.1492, p=6.3 x 10–07); angiogenesis (r=0.3059, p=6.8 x 10–25); and hypoxia based on Elvidge et al (r=0.5320, p=1.2 x 10–79) and negative correlation with hormone expression (Fig S2a, 2b and 2c). Consistent with this, *CRYAB* overexpression predominantly occurs in triple-negative breast cancer that lack expression of hormone receptors^7^. Notably, autophagy, angiogenesis and, hypoxia have a significant-positive correlation to *CRYAB* expression across several cancer types such as adenocarcinomas of colon, lung, prostate, and stomach (Fig S2c,). In the clinical setting, autophagy and angiogenesis are linked to *CRYAB* overexpression and this gene is abundantly expressed in hypoxic regions of tumors; however, in cell-based *in vitro* assays the reoxygenation and subsequent generation of reactive oxygen species (ROS) activates *CRYAB* rather than hypoxia^2^.

Next, to investigate correlation between gene expression data and estimation of the abundances of immune cell types, we conducted a Cibersort query which revealed a positive correlation between *CRYAB* expression and macrophage infiltration, particularly M2 macrophages, which play a significant immunosuppressive role in tumor microenvironment in bladder (r=0.3954, p=2.6e–16, n=397), colon (r=0.4032, p=1.1e–18, n=442), esophageal (r=0.3692, p=5.3e-07, n=174), head and neck squamous cell carcinoma (r = 0.1984, p = 6e-06, n =516), kidney (r = 0.2790, p = 2.2e-06, n = 279), liver (r = 0.1832, p = 0.0005, n = 362), rectum (r = 0.5579, p = 3.8e–14, n = 156) and, testicular germ cell tumors (r = 0.2269, p = 0.0054, n = 149). However, a negative correlation between *CRYAB* expression and macrophages was found in breast (r = −0.1526, p = 4.2e-07, n = 1090), prostate (r = −0.1940, p = 8.4e-05, n = 406) and sarcoma (r = 0.1945, p = 0.0035, n = 224) (Fig S2d). Additionally, we found negative correlation of *CRYAB* expression and lymphocytes in a large number of tumor types, suggesting reduced lymphocytic infiltration. Overall, the bioinformatic analysis suggests that *CRYAB* overexpression leads to poor outcome in several types of cancers and angiogenesis is significantly positively correlated to *CRYAB* expression in most cancers.

### *Cryab* overexpression in the mouse model causes spontaneous tumorigenesis *in vivo*

Both mouse and human αB-Crystallin proteins are comprised of 175 amino acids and exhibit 98% sequence homology with just 4 amino acid mismatches^9^. Between these two species, the three sites of phosphorylation and the α-B crystallin domains are conserved (Fig S3a). Collectively, αB-Crystallin is a conserved protein in human and mouse. To characterize the pathophysiological role of *Cryab* overexpression *in vivo*, we generated a Flag-*Cryab* knock-in mouse model to ubiquitously overexpress *Cryab* cDNA from the Rosa26 locus (Rosa26 Tg/Tg) (Fig 1a). Correct targeting was validated by PCR on genomic DNA extracted from different genotypes (*Cryab*^Wt/Wt^ = Wt, *Cryab*^Wt/Tg^ = Het, *Cryab*^Tg/Tg^ = Tg or Hom) (Fig S3b) and protein expression across different organs of the mouse using immunoblotting analysis, respectively (Fig S3c). *Cryab*^Tg^ mice were viable and had no obvious developmental defects. Using this constitutive overexpression mouse model of *Cryab*, we monitored a cohort of wild type mice (hereafter referred to as *Cryab*^Wt^, n=40) as a control group to compare with homozygous transgenic mice overexpressing αB-Crystallin (*Cryab*^Tg^, n = 40: 23 Females, 17 Males) over time for spontaneous tumor formation. Any mouse showing signs of distress and disease according to the ethical limitations was sacrificed for further histopathological investigation and considered as the experimental endpoint.

**Fig 1.**
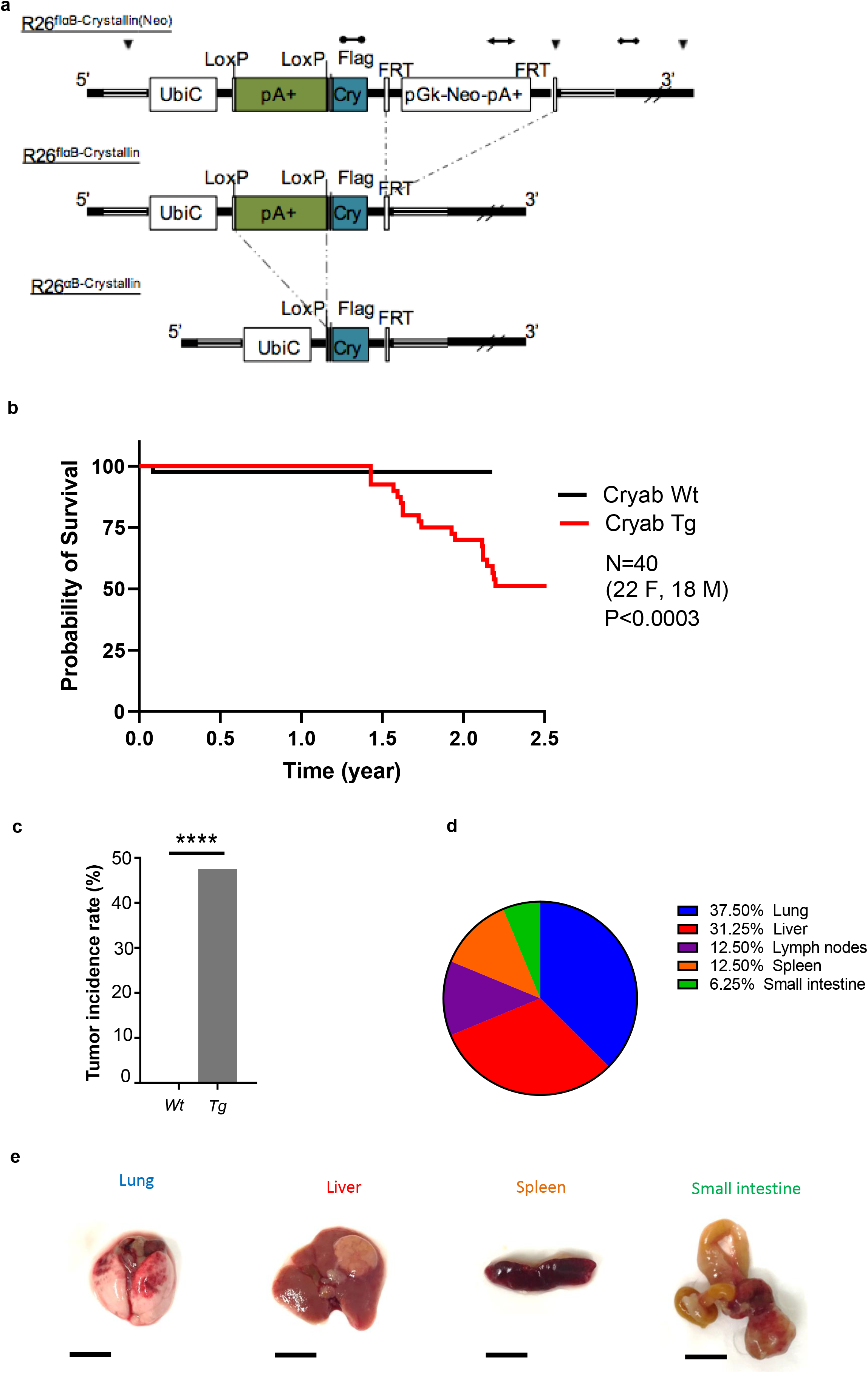
*Cryab* overexpression causes spontaneous tumorigenesis in vivo. **(a)** Schematic representation of αB-Crystallin overexpressing mice (Ozgene) developed by knocking in the transgene into the Rosa26 locus. The targeting vectors show recombination steps. Legend: FLPe recombination, Cre recombination, Homology arm, ▼ EcoRV site, transgene probe (tgP), 3’ probe (3’P), Neomycin probe (Neo). **(b)** Kaplan–Meier survival analysis of mice for indicated genotypes (n = 40 per group) Log-rank (Mantel–Cox) test was performed to determine P-value < 0.00001. **(c)** Percentage of cancer incidence rate among mice of indicated genotypes (n ▾ 40 per group); Fischer exact test was performed to determine P-value < 0.00001 (****). **(d)** The distribution of tumor spectrum in tissues of *Cryab* transgenic mice. **(e)** Representative images of gross morphology of tissues with different malignancies. (Scale bars, 1 cm)

*Cryab*^Tg^ were more susceptible to the formation of tumors compared to the control group as shown in Kaplan-Meier tumor-free survival analysis (Fig 1b). The spontaneous tumorigenesis in *Cryab*^19^ started at 17 months and by 26 months half of these mice died due to tumor burden of at least one tumor type while in this time frame just one mouse from the control group died due to an idiopathic cause. While the tumor incidence was 50% during the period of monitoring (Fig 1c), all mice were subjected to examination for signs of macroscopic tumor formation across major organs and tissues including mammary glands, liver, lung, spleen, small intestine, kidney and lymphoid. The major types of tumors observed originated from lung and liver followed by lymph nodes, spleen and small intestine (Fig 1d, 1e, Table 1). The pathological investigation of tumorigenesis based on hematoxylin and eosin (H&E) staining revealed further details of the tumor spectrum including alveolar/bronchiolar adenocarcinomas, hepatocellular carcinomas, B-cell lymphomas, sarcomas as well as liver and lung metastases (Fig 2a-2g, Table 1). Collectively, these data highlight that *Cryab* overexpression is sufficient to cause a broad spectrum of spontaneous primary and metastatic tumors in mice with 50% incidence in the time-frame of this study.

**Table 1.** The distribution of cancer spectrum in Cryab transgenic mice.

**Fig 2.**
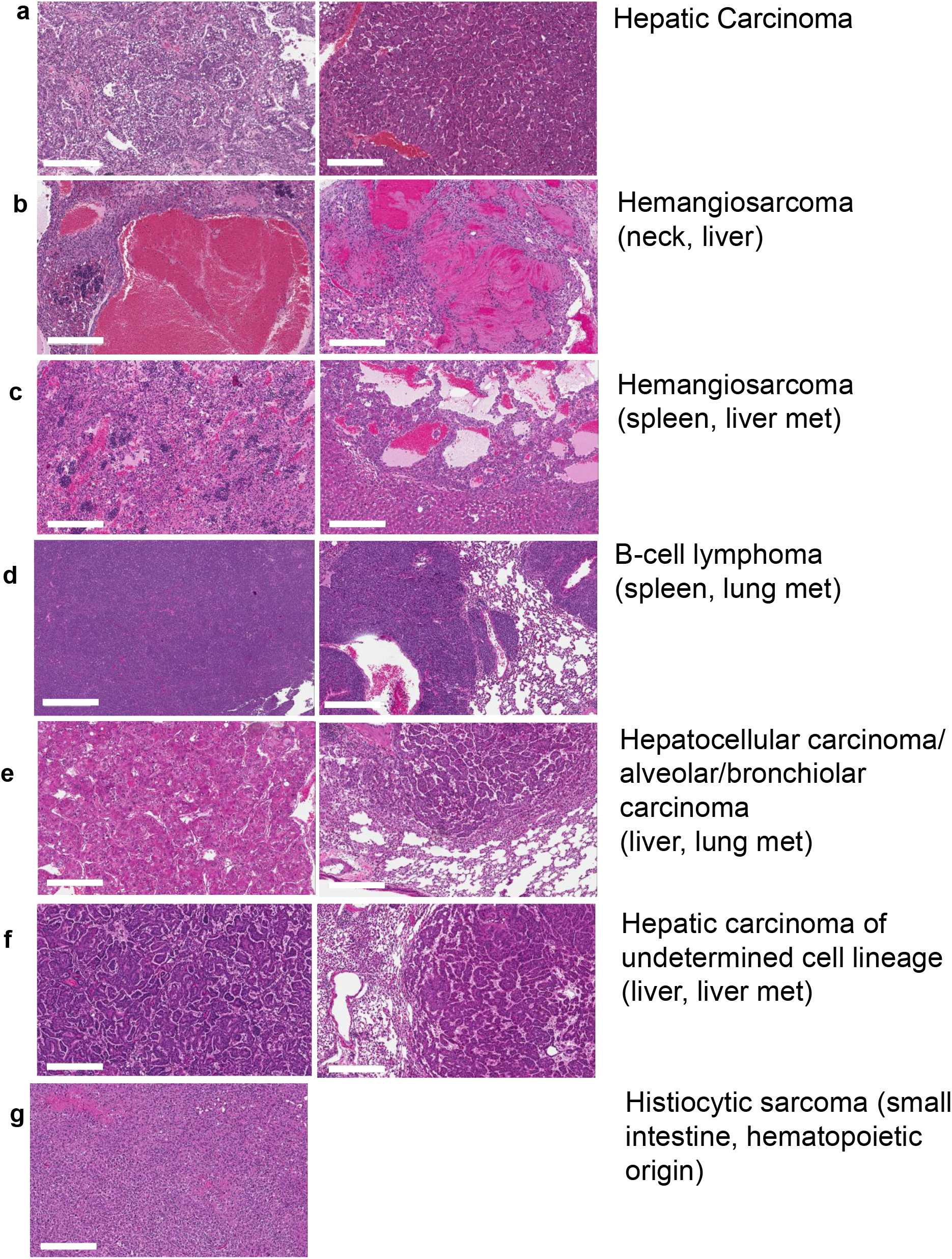
Characterizing and classifying the primary tumors and metastasis. **(a-g)** Representative H&E stained microscopic images of selected sections of tumor-bearing *Cryab*^Tg^ mice as indicated in the figure (scale bars, 300 μm).

Given that constitutive overexpression of *Cryab* leads to malignancies with late latency ~17 months, next we sought to validate this observation by using a 7,12-Dimethylbenz[a]anthracene (hereafter: DMBA)-induced carcinogen model to evaluate differences in the tumorigenic potential of *Cryab*^Wt^, *Cryab*^Wt/Tg^ and *Cryab*^Tg^ mice. DMBA is a potent organ-specific carcinogen that serves as a tumor initiator. At postnatal day 5, pups from each genotype were subjected to carcinogen-induced tumorigenesis by topical administration of 1% DMBA^10^ and mice were sacrificed at 4 months and analyzed for the development of lung tumors. We found that *Cryab*^Tg^ exhibited a higher number of lung tumor nodules when compared to *Cryab*^Wt^ and *Cryab*^Wt/Tg^ counterparts suggesting that *Cryab* overexpression can potentiate carcinogen-induced tumorigenesis (Fig S4a, S4b).

### *Cryab* overexpression is correlated with increased angiogenesis and epithelial to mesenchymal transition in tumors

Given the spontaneous formation of tumors and associated metastases in *Cryab*^Tg^ mice; first, we validated the expression level of αB-Crystallin in *Cryab*^Wt^ and *Cryab*^Tg^ tissues. Upon immunohistochemistry staining of αB-Crystallin, we found that it was elevated significantly in liver, spleen, and lung tumors from *Cryab*^Tg^ mice when compared to respective age-matched tissues from *Cryab*^Wt^ mice (Fig 3a, 3b). Next, we examined the level of immunohistochemical expression of well documented cancer relevant genes and pathways including TP53, extracellular signal-regulated kinase (ERK) and Protein Kinase B (AKT) in matched tissues of normal and cancer-bearing mice. Expectedly, we observed a higher expression of these markers in tumors from *Cryab*^Tg^ mice compared to age-matched control tissues from *Cryab*^Wt^ (Fig S5a). Consistently, we found that cancers in *Cryab*^Tg^ mice had higher levels of p53 when compared to adjacent normal tissues and/or corresponding tissues from *Cryab*^Wt^ mice (Fig S5a). The observed increase in p53 expression is most likely an indication of the presence of mutated p53 which might act as a secondary hit to initiate tumorigenesis in *Cryab*^Tg^ mice. Both oncogenic survival signaling pathways, ERK and AKT, might also be activated by *Cryab* in response to stress-induced mechanisms in tumor microenvironment (Fig S5a).

**Fig 3.**
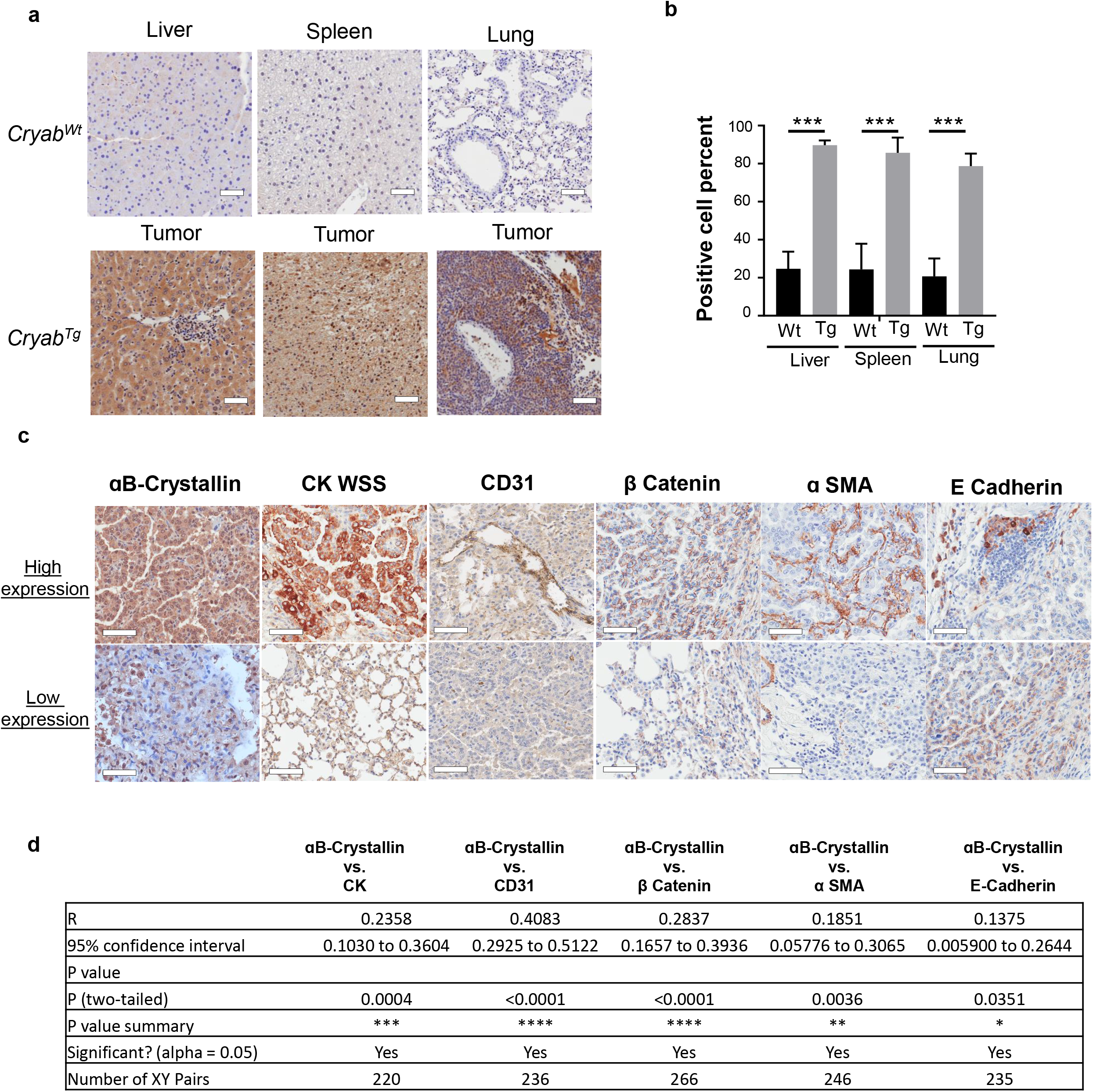
Transgenic expression of *Cryab* correlates with progressive tumorigenesis and tumor markers. **(a)** Representative microscopic images of normal organs in *Cryab*^wt^ (upper Panel) and *Cryab*^Tg^ (lower Panel) associated malignancy stained with anti-Crystallin antibody (scale bars, 50 μm) **(b)** Bar chart represents the positive cell count percent of selected tissues for indicated genotypes. **(c)** Comparison of high and low expression IHC staining of indicated markers in adenocarcinoma (ADC) of lung of tumor-bearing *Cryab*^Tg^ mice. **(d)** Correlation comparison between indicated markers shown above.

Next, we examined whether there is a correlation between the pattern of *Cryab* expression in tumors from *Cryab*^Tg^ mice and the expression of tumor markers in malignant tissues. Tumor types were divided into lymphoma, hepatocellular carcinoma, lung and, liver adenocarcinoma and regions of interest were highlighted based on cytokeratin-wide spectrum screening (CK-WSS) and gross morphology of tumors. The respective histoscore (H-score) for tumor and tumor-adjacent tissues were obtained according to previously published method explained in method section^11^. Interestingly, there was a strong correlation between cytokeration-wide spectrum screening CK-WSS, a marker of epithelial cancer cells and αB-Crystallin expression in solid tumors (Fig 3c, 3d and Supplementary table 1). Furthermore, we found a positive correlation between αB-Crystallin and CD31, a marker of angiogenesis in tumors. There was also a positive correlation between αB-Crystallin and membranous β Catenin which is associated with various carcinomas and epithelial-to-mesenchymal transition (EMT). Moreover, the attenuation of E-cadherin compared to stronger expression of α-smooth muscle actin (SMA) is also linked to EMT. However, we could not find a significant immunohistochemical correlation between αB-Crystallin expression and Ki67 proliferation marker nor matrix metalloproteinase-9 (MMP-9) expression in lung and liver adenocarcinomas (Fig S5b). These markers are relatively high in fast-dividing proliferative cells and in advanced stage aggressive cancers and metastasis. On the contrary, Ki67 in lymphoma and MMP9 in hepatocellular carcinoma was comparatively increased (Fig S5b). Taken together, we found a causal link between αB-Crystallin expression and crucial events of tumor initiation and spread in particular angiogenesis and EMT, in agreement with bioinformatic analysis of human tumors shown in (Fig S2a and S2C).

### *Cryab* overexpression promotes colony formation, migration and survival signaling *in vitro*

The identification that *Cryab* overexpression causes spontaneous tumor formation in multiple organs and is linked to essential oncogenic markers *in vivo* prompted us to evaluate the molecular mechanism underpinning these phenotypes. Consequently, mouse embryonic fibroblasts (MEFs) from *Cryab*^Wt^, and *Cryab*^Tg^ transgenic mice (Fig S6a) were generated and transformed using a construct expressing E1A/RasV12 oncogenes^12^ to evaluate signaling roles of αB-Crystallin. Heat-shock stress (43°C) was used to monitor the induction of *Cryab* expression at various time-points (Fig S6b). As *Cryab*^13^ MEFs demonstrated significantly higher expression of αB-Crystallin compared to *Cryab*^Wt^ MEFs within 1 hour after heat-shock stress, this time-point was utilized for subsequent experiments. Next, we compared the proliferation potential of MEFs using IncuCyte^®^ live cell analysis for 6 days. Despite marginally increased growth in one of the *Cryab*^Tg^ MEF lines, no significant differences between the proliferation rate of *Cryab*^Wt^ and *Cryab*^Tg^ MEFs were observed (Fig 4a). However, *Cryab*^13^ MEFs had enhanced clonogenic potential as assessed by colony-forming ability compared to *Cryab*^Wt^ counterparts (Fig 4b) and the differences in clonogenic survival were accentuated after heat-shock treatment (Fig 4c). Moreover, the transwell migration assay demonstrated that *Cryab*^Tg^ MEFs showed increased migratory potential compared to *Cryab*^Wt^ MEFs following heat-shock treatment (Fig 4d).

**Fig 4.**
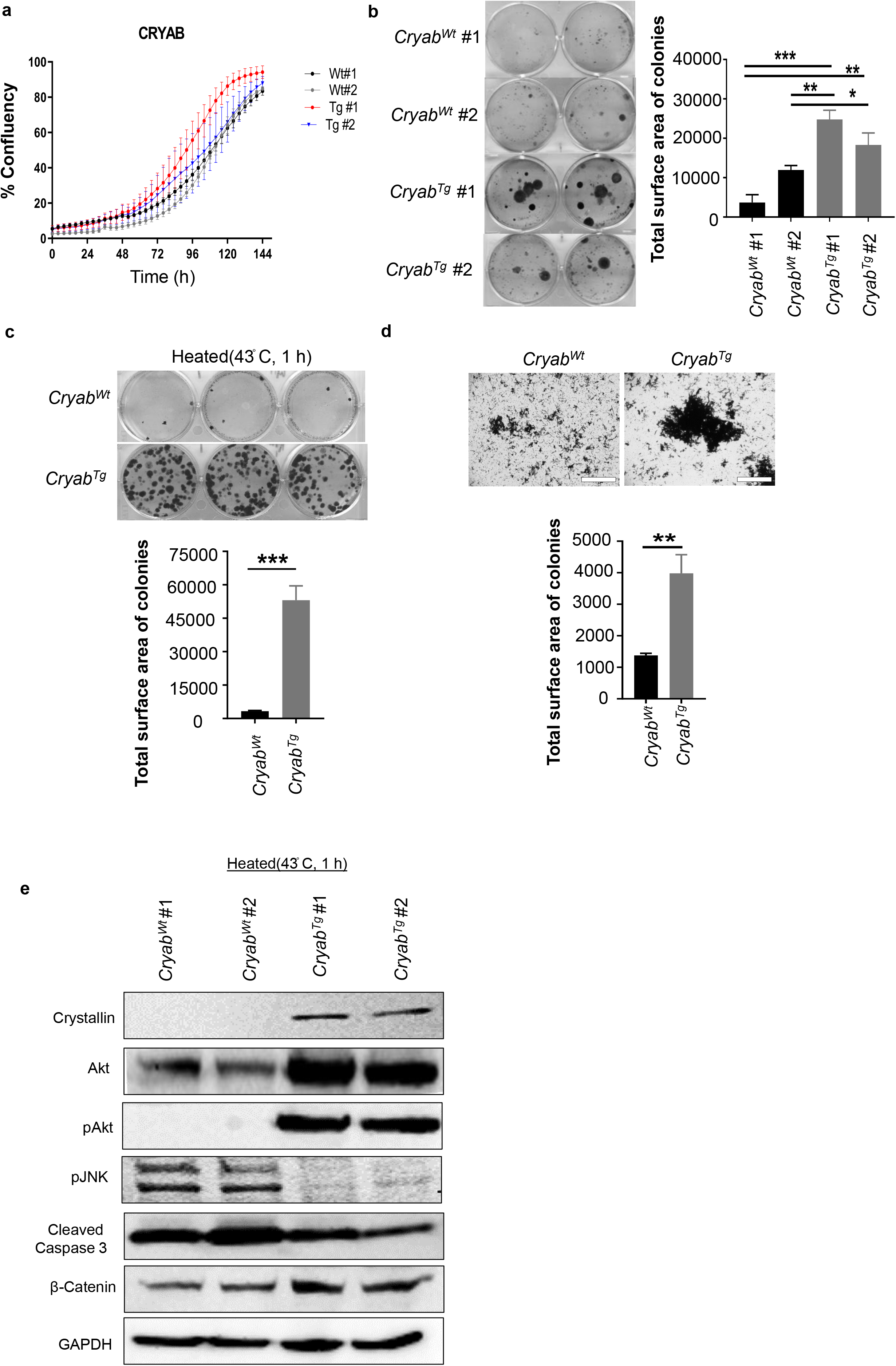
*Cryab* overexpression increases clonogenic survival and transwell migration in vitro. **(a)** Proliferation of E1A/Ras-transformed MEFs *Cryab*^Wt^ (Wt) and *Cryab*^Tg^ (Tg) measured using IncuCyte (Mean ± SEM, an average of 2 biological repeats in duplicate and 2 independent experiments Student’s t-test, ****P < 0.0001), **(b)** Representative image (left) and quantification (right) of indicated genotypes of E1A-Ras-transformed MEFs for clonogenic survival (colony formation) assay. Each cell line was seeded at densities of 500 in each well incubated for 14 days, the cells were then processed using the crystal violet stain to visualize the colonies. Quantification performed by analysis of all images **(c)** Representative image (upper) and quantification (lower) of indicated genotypes of E1ARas-transformed MEFs for clonogenic survival (colony formation) assay as aforementioned above after initial 1 hour of heat stress (43°C). **(d)** Representative image (upper) and quantification (lower) of wells for indicated genotypes of E1ARas-transformed MEFs for transwell migration assay after an initial 1 hour of heat stress (43°C), (scale bars, 400 μm) Data in (b-d) presented as Mean ± SEM, an average of 3 biological repeats and 3 independent experiments Student’s t-test, ****P < 0.0001). **(e)** Immunoblot analysis was performed on E1A/Ras-transformed MEFs after exposure to heat stress at 43°C for 1 hour.

Next, we investigated possible signaling pathways altered by the overexpressed αB-Crystallin in MEFs. We found that the levels of total AKT and phosphorylated AKT at residue 473 (pAKT 473) were both upregulated (Fig 4e). The PI3K/AKT is a major survival pathway that plays multiple roles in cellular processes such as regulation of cell proliferation, apoptosis, metabolism, and cell migration. To further examine the role of αB-Crystallin in apoptosis evasion, we examined the induction of apoptotic pathways following heat shock. We found that the expression of the pro-apoptotic phosphorylated JNK (pJNK T184/Y185) and apoptotic marker cleaved caspase-3 were both downregulated in the *Cryab*^Tg^ lines (Fig 4e). Importantly, β Catenin had elevated expression in *Cryab*^Tg^ MEFs which is consistent with IHC staining in tumors which indicated the activation of the Wnt signaling pathway. Overall, *Cryab* overexpression promotes clonogenic potential, increases cell migration and cell survival signaling and inhibits apoptosis.

### *Cryab* overexpression upregulates metastatic and oncogenic markers

To investigate the signatures enhancing the oncogenic potential of *Cryab*^Tg^ MEFs, we evaluated the proteome landscape of *Cryab*^Wt^ and *Cryab*^Tg^ MEFs. The mass spectrometry-based proteomic analysis revealed 135 downregulated and 104 upregulated proteins >log_2_ (±0.6) in *Cryab*^Tg^ MEFs over *Cryab*^Wt^ after 1 hour heat-shock (43^3^ C) stimulation. Gene enrichment analysis identified differential regulation in biological pathways such as epithelial morphogenesis/migration (migration), developmental growth, hypoxia, mitotic cell cycle, response to leukocytes (activation and proliferation in immune system process), regulation of apoptosis, extracellular remodeling and wnt signaling in *Cryab*^Tg^ MEFs (Fig 5a). The volcano plot of downregulated and upregulated proteins revealed the plasma membrane immunoglobulin-like protein Basigin (BSG, CD147), gained the highest fold change and p value among the others (Fig 5b). Basigin is overexpressed in many different human cancer types and plays a crucial role in regulation of cancer cell invasion, migration and angiogenesis^13^. Basigin is strongly connected to protein clusters for epithelial morphogenesis/migration, developmental growth and immunity. Basigin with linked upregulated proteins enhancing metabolism and growth such as growth factor HBEGF, glucose transporter SLC2A1 and proto-oncogene JUNB, combined with downregulated protective pathways containing donor against oxidative stress G6PDX, redox chaperone PARK7 (purine metabolism) and DNA repair enzyme APEX1 (catabolic nucleobase compounds)^13^, likely provide an advantage for overall oncogenic growth and transformation (Fig 5a, Fig S7a). Supporting the biological pathway enrichment analysis, the gene annotation of upregulated genes linked most of them to metastasis and tumorigenic phenotypes (Supplementary table 2). Intriguingly, top five upregulated genes are all linked to regulation of oncogenesis and metastasis (Fig S7b) and pan cancer analysis of TCGA dataset revealed that these genes have co-occurrence of expression in cancers including CRYAB with BSG and HELLS (Fig S7c). Further supporting relevance to human cancer development, CRYAB expression levels are significantly higher in metastatic contexts in a range cancer types (Fig S8). Additionally, CRYAB expression significantly correlates with EMT in various cancer types and with angiogenesis in 14 of 22 (64%) cancer types of epithelial origin as already shown in Fig S2.

**Fig 5.**
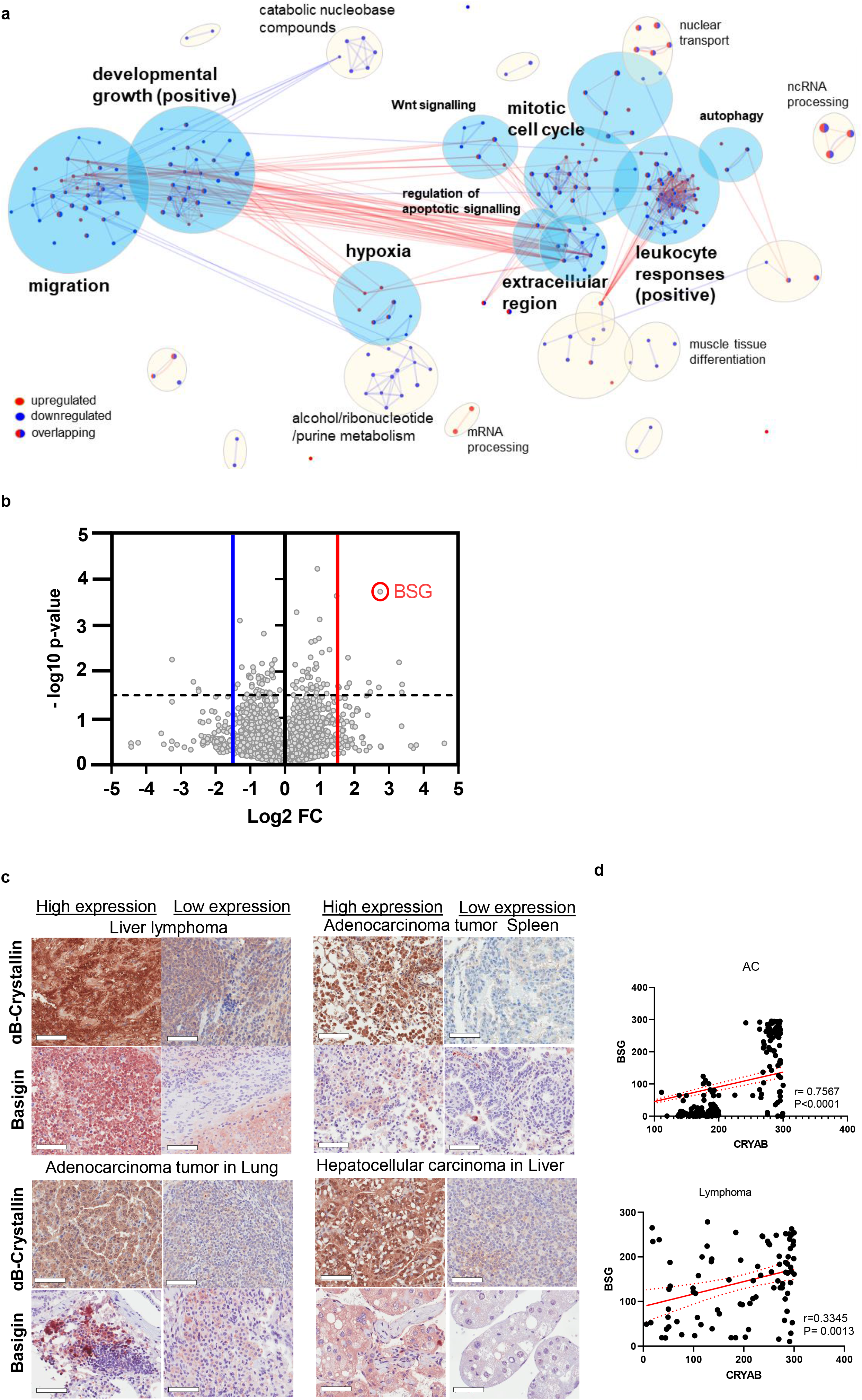
Mass spectrometry (MS)-based proteomics analysis. **(a)** Biological pathway enrichment analysis of the proteomic data comparing Cryab Tg E1ARas-transformed MEFs to Cryab WT MEFs revealed differentially regulated pathways visualized using Cytoscape. Upregulated (red) and downregulated (blue) node proteins in clusters are indicated. **(b)** Volcano plot showing up/downregulated proteins (*Cryab*^Tg^/WT) in the proteomic data with corresponding fold changes and p-values. Highlighted BSG had the highest p-value with ~7-fold upregulation. **(c)** Representative IHC images for comparison of Cryab (high and low expressing areas) and BSG expression in Lymphoma (Lym) liver, adenocarcinoma (ADC) of lung, liver and hepatocellular carcinoma (HCC) of tumour-bearing *Cryab*^Tg^ mice, (scale bars, 50 μm). **(d)** Correlation (linear regression) between expression of Cryab and Bsg in adenocarcinoma and lymphoma of tumour-bearing CryabTg mice.

To validate these finding, we performed immunostaining of Basigin in tumor tissues of *Cryab*^Tg^ mouse model. These data validated that Basigin protein expression positively correlates with *Cryab* expression in tumors and adjacent tumor tissues (Fig 5c, 5d). Next, we performed the immunofluorescence staining on MEFs and the data validated the higher expression of *Cryab* (*red*) and BSG (green) at the plasma membrane in *Cryab*^Tg^ MEFs, both of which decreased upon BSG knockdown by siRNA (Fig 6a). Interestingly, BSG knockdown also led to the reduction of colony formation in *Cryab*^Tg^ MEFs (Fig 6b). Furthermore, BSG knockdown significantly decreased migration in *Cryab*^Wt^ and *Cryab*^Tg^ MEFs by transwell migration assay (Fig 6c). Collectively, the data suggest that some of the phenotypic effects of *Cryab* overexpression might-in part be mediated by BSG expression in our murine and cellular models.

**Fig 6.**
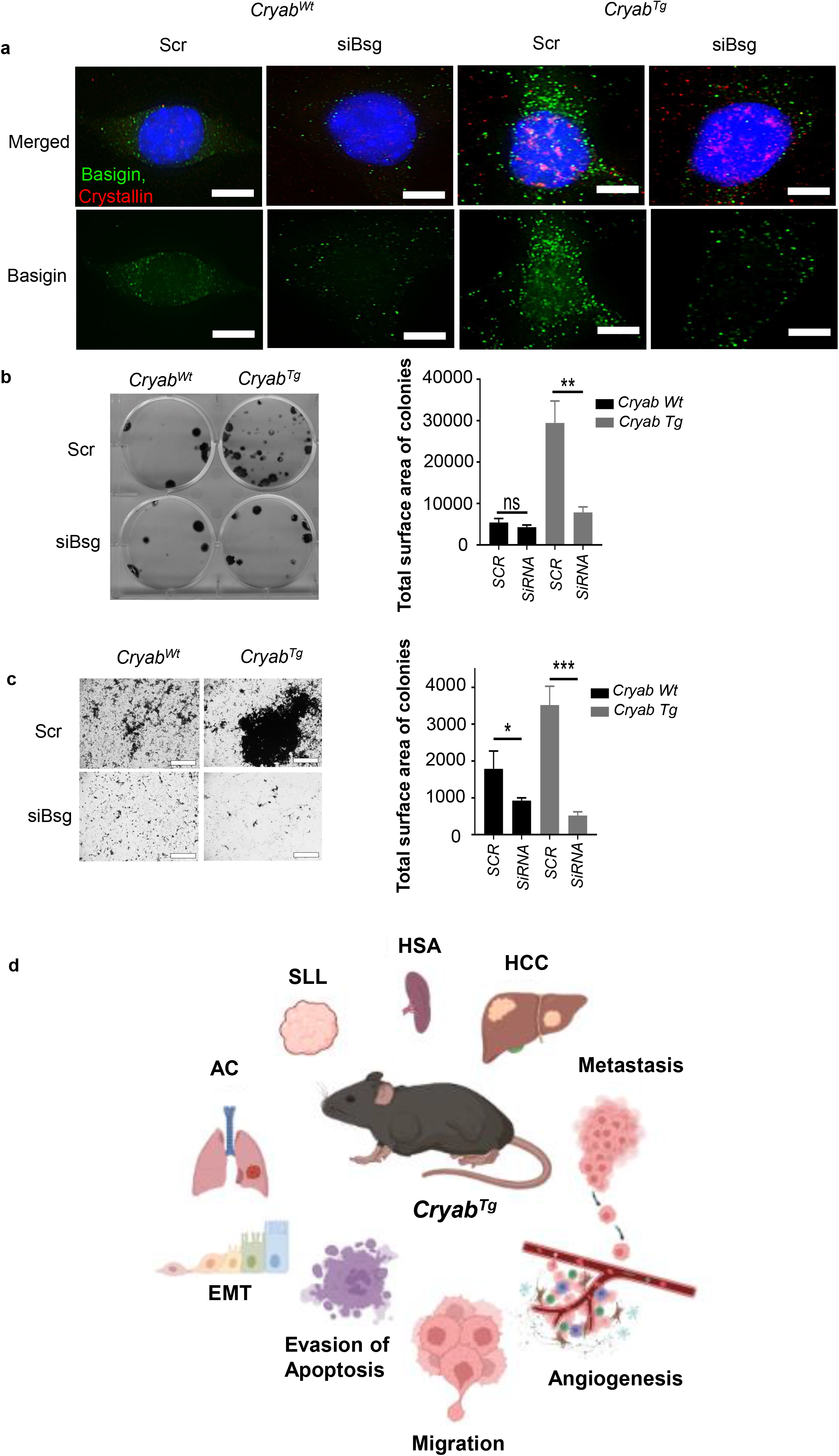
Basigin (BSG) inhibition reduces colony formation. **(a)** Representative image of *Cryab*^Wt^ and *Cryab*^19^ of E1ARas-transformed MEFs treated with Scr (scrambled) and siRNA against *Bsg* showing the expression of *Cryab* (red) and BSG (green). (scale bars, 15 μm) **(b,c)** representative image (upper) and corresponding quantification of **(b)** colony formation assay **(c)** transwell migration assay (lower) for Cryab wt and Cryab Tg of E1ARas-transformed MEFs treated with Scr (scrambled) and siRNA against BSG, (scale bars, 400 μm), Data presented as Mean ± SEM, an average of 3 biological repeats and 3 independent experiments Student’s t-test, ****P < 0.0001). **(d)** The graphical abstract of *Cryab*^Tg^ mouse model shows spontaneous tumorigenesis and metastasis *in vivo* leading to different cancer types such as Hepatocellular carcinoma (HCC), Hemangiosarcoma (HSA), Small lymphocytic lymphoma (SLL), and Adenocarcinoma (AC) due to regulation of several pro-tumorigenic mechanisms including EMT, evasion of apoptosis, angiogenesis and migration.

In conclusion, we have characterized the mouse model of *Cryab*^13^ which led to the spontaneous formation of tumors and associated metastasis due to pro-tumorigenic alteration in promoting survival signalling and migration-EMT and evasion of apoptosis (Fig 6d).

## Discussion

To understand the contribution of αB-Crystallin to *de novo* tumorigenesis, we generated and characterized a novel mouse model with transgenic overexpression of *Cryab*. Previous studies using human cancer cell lines and human xenograft models have reported both pro- and anti-tumorigenic role of αB-Crystallin^6^. This transgenic mouse model is the first, to the best of our knowledge, to show that *Cryab* overexpression is sufficient to induce tumorigenesis *in vivo*. Mice with homozygous *Cryab*^Tg^ expression developed a wide spectrum of solid and hematological tumors with an approximate 50% incidence rate and metastatic potential, although with late latency. These findings provide evidence to support that *Cryab* can act as an oncogene. Strikingly, we found higher p53 protein levels, which are most likely an indication of *Trp53* mutation in representative *Cryab*^Tg^ tumor tissues, suggesting that Tp53 might be a critical secondary hit required for tumor initiation observed in the homozygous *Cryab*^Tg^ mice. The tumors were further characterized histopathologically which identified lung adenocarcinoma, lung metastasis, hepatocellular carcinoma, and lymphoma as the major types of malignancies. Furthermore, *Cryab*^Tg^ mice also showed the increased carcinogen-DMBA induced tumor load. Like *Cryab* (also known as *HSPB5*), other heat shock proteins have been functionally linked to spontaneous tumorigenesis, for example, transgenic mice expressing human HSP70 developed lung and lymph node tumours before 18 months of age. In addition, HSP27 increased the tumorigenicity of rat colon adenocarcinoma in a syngeneic model^14^. Furthermore, HSP90 is similarly up-regulated in a wide variety of cancers and inhibitors of HSP90 are currently in clinical trials as chemotherapeutic drugs^15^.

In our transgenic mouse model, we found elevated levels of *Cryab* expression in *Cryab*^Tg^ tumors compared to adjacent non-tumor tissues and age-matched normal tissues from *Cryab*^Wt 2^. *Cryab* is a multi-functional protein with postulated roles in the regulation of cell architecture, apoptosis, and autophagy through interactions with a multitude of its substrate proteins^16^. Therefore, there are several potential mechanisms by which *Cryab* may promote tumorigenesis. We have found that this may be caused by elevated levels of AKT and ERK survival pathways in *Cryab*^Tg^ tissues (both tumors and adjacent tissues) compared to tissues from *Cryab*^Wt^ mice. Interestingly, *Cryab* overexpression was correlated with cytokeratin-screening (CK-WSS), which reflects tumor cell activity and is considered a tumor marker. Furthermore, we found a correlation between overexpressed *Cryab* and both angiogenesis and EMT markers in tumors, which may both enhance tumor phenotypes. The negative consequence of *Cryab* depletion on tumor angiogenesis has been established in *in vitro* studies using cancer cell lines although, increased *Cryab* expression has not previously been functionally linked to increased angiogenesis^17^’^18^. Here, we find that high *CRYAB* expression is also linked to angiogenesis, EMT and metastasis in a variety of human cancer types, suggesting that our observations are relevant to cancer development in patients.

To complement and extend prior knowledge of αB-Crystallin biology, we performed further phenotypic analysis of E1A/RAS transformed MEFs derived from *Cryab*^Tg^ and *Cryab*^Wt^ mice. Notably, we show that overexpression of *Cryab* in transformed MEFs is sufficient to impart tumorigenic properties including increased clonogenic capacity, and migratory and invasive potential. Functional consequences of *Cryab* overexpression were examined by proteomics analysis of the transformed MEFs of both genotypes, which also pointed to pleiotropic roles of *Cryab* in the regulation of many metastatic and oncogenic proteins. The most significantly upregulated protein was Basigin, which we found had a positive correlation with the expression of *Cryab*, validated both in tumors compared to adjacent normal tissues and *in vitro* in MEFs. Basigin is also a transmembrane protein that mainly functions in metabolic pathways, such as glycolysis, but its overexpression is associated with several pathologies including cancer^19–22^. In particular, its overexpression is correlated with worse overall survival in acute myeloid leukaemia, and non-small-cell lung cancer (NSCLC) and plays important role in regulation of cancer cell proliferation, invasion and metastasis^20,22,23^. Its cancer connection is mostly linked to its capacity to regulate expression/activity of monocarboxylate transporters, matrix metalloproteinases and PI3K and MAPK pathways. It also functions as a key mediator of inflammatory/immune response. Interestingly, the knockdown of Basigin reduced the oncogenic colony formation ability and migratory potential of *Cryab*^Tg^ MEFs, suggesting that some of the phenotypic effects of *Cryab* overexpression might-in part be mediated by regulation of Basigin expression.

We also found elevated levels of AKT and pAKT in *Cryab*^Tg^ MEFs, consistent with our observations in *Cryab*^Tg^ tumors, and this might be a potential cause of spontaneous tumorigenesis in our model. The PI3K/AKT pathway is involved in various cellular processes, including the promotion of cell survival, and cell cycle and is often altered in various cancers^24^. AKT can inhibit apoptosis by phosphorylating and inhibiting pro-apoptotic proteins such as Bad, Bim and caspase 9^25^. It can also promote the cell cycle progression by inhibiting both Cyclin-Dependent Kinase (CDK) degradation and CDK Inhibitors expression, allowing cell cycle activation^24^. The upregulation of pAKT and total AKT by *Cryab* overexpression are consistent with another *in vitro* model in particular near-normal mammary epithelial cell line, MCF10A, in which *Cryab*-overexpression promoted malignant transformation and growth of mouse mammary xenograft tumors through regulation of AKT^8^. On the other hand, JNK was found to be downregulated in *Cryab*^Tg^ MEFs. JNK directly induces apoptosis by activating pro-apoptotic proteins such as the previously mentioned Bad and Bim and inhibiting anti-apoptotic proteins such as Bcl-xL. JNK has been reported to antagonize PI3K/AKT pathway, and the reduced apoptosis observed in our model may partly potentiate the tumor formation where JNK antagonism of Akt-mediated survival signals is suppressed. Evasion of apoptosis is a well-described hallmark of cancer resulting in cellular resistance to conventional chemotherapeutic agents^1^. Indeed, resistance to apoptosis during myocardial ischemia was observed in mice overexpressing *Cryab* in cardiomyocytes^26^. Altogether, these results show that the overexpression of CRYAB leads to the promotion of cell survival after induction of cellular stress, which is consistent with its oncogenic potential. This may occur via the activation of the AKT pathway and inhibition of the JNK pathway. The promotion of cell survival was validated by the decreased levels of cleaved caspase 3.

In summary, this is the first report to our knowledge that has causally linked *Cryab* to oncogenesis. Our data demonstrate that *Cryab* overexpression beyond a critical level is self-sufficient to induce a wide spectrum of spontaneous tumors in aged mice. We have found that this may be caused by an upregulation of the oncogenic AKT survival pathways, and modulation of the tumor suppressor JNK pathway. Notably, overexpression of *Cryab* in transformed MEFs is sufficient to impart tumorigenic properties and further studies are required to enhance the knowledge of how *Cryab* regulates these phenotypes, especially in a hypoxic and oxidative stress environment of tumor cells. Our mouse model could be a valuable tool in studying the mechanism of oncogene-related tumorigenesis, in particular given the consistency between our observations in mice and patients. The crossing of *Cryab* mouse line with other cancer-prone models such as KrasG12D expression, PTEN and/or p53 loss will highlight the contribution of *Cryab* to cancer progression and its potential as a future treatment target.

## Materials and Methods

### Generation of the targeting construct

Generation of Rosa26-UbiC-Cryab floxed mice was a contracted service performed by Ozgene (Perth, WA, Australia). To generate the *Cryab* knock-in model, we designed a targeting vector composed of Flag-tagged Cryab cDNA (fl-Tg) preceded by a human ubiquitin C (UbiC) promoter and lox-P flanked polyadenylation (pA+) stop region, with a downstream flippase recombinase target site-flanked neomycin resistance cassette (PGK-NEO) for embryonic stem cell (ESC) selection. Genomic targeting of the construct was achieved in wild-type BALB/C ESCs, by using standard homologous recombination and blastocyst manipulation techniques. Gene manipulation was verified by Southern blot analysis by Ozgene, with probes against both the endogenous coding region and NEO selection cassette following restriction digest of genomic DNA with the EcoRV restriction enzyme.

### Generation of the ubiquitous Cryab knock-in mouse model

Cryab knockin mice were generated by crossing heterozygous Rosa26Ubiq-polyA-flTg(New)/wt mice with FLPe mice to excise the PGK-Neo cassette, followed by backcrossing to C57BL/6 wild-type mice to remove the FLPe transgene. Rosa26UbiC-polyA-flTg(Neo)/wt mice were subsequently crossed with Rosa26EIIA-Cre mice (Ozgene) to excise the (pA+) stop region, allowing overexpression of Cryab cDNA from the Rosa26 locus (Rosa26Tg/Tg). These mice were then crossed to BALB/C wild-type mice to segregate the Rosa26EIIA-Cre and RosT26Tg/Tg alleles. The resulting Cryab heterozygous (Cryab^Wt/Tg^) mice were intercrossed to generate 3 genotypes: wild-type (Cryab^wt/wt^), heterozygous (Cryab^wt/Tg^), and homozygous (Cryab^Tg/Tg^) mice. Wild-type control mice used for Kaplan-Meier survival analysis were of a mixed C57Bl/6/Balb/C genetic background and served as joint controls with^27^.

### Animal husbandry and ethics statement

All experimental animals were maintained on a mixed background (Balb/c X C57BL/6) strain and were housed at 25°C in a 12 h light-dark cycle. This study was performed in strict accordance with the Australian code for the care and use of animals for scientific purposes. This work was approved by the Queensland Institute of Medical Research (QIMR) Berghofer Medical Research Animal Ethics Committee (number A0707-606M).

### Cell culture

Mouse embryonic fibroblasts were generated as described previously^28^. Briefly, primary MEFs were immortalized by E1A/Ras transduction. All cell lines were annually authenticated using short tandem repeat (STR) profiling and routinely tested for Mycoplasma infection by scientific services at QIMR Berghofer Medical Research Institute.

The cells were maintained in culture in Dulbecco’s Modified Eagle’s Medium (DMEM) (Life Technologies TM, Carlsbad, CA, USA) containing 20% Fetal Bovine Serum (SAFC BiosciencesTM, Lenexa, USA) 1% penicillin-streptomycin (Life Technology TM) and 1% Amphotericin B

### siRNA Transfection

MEFs were plated in 6 well plates at density of 200,000 cells/well followed by double reverse transfection using 20 nM of siRNAs with following sequence (siRNA against BSG):

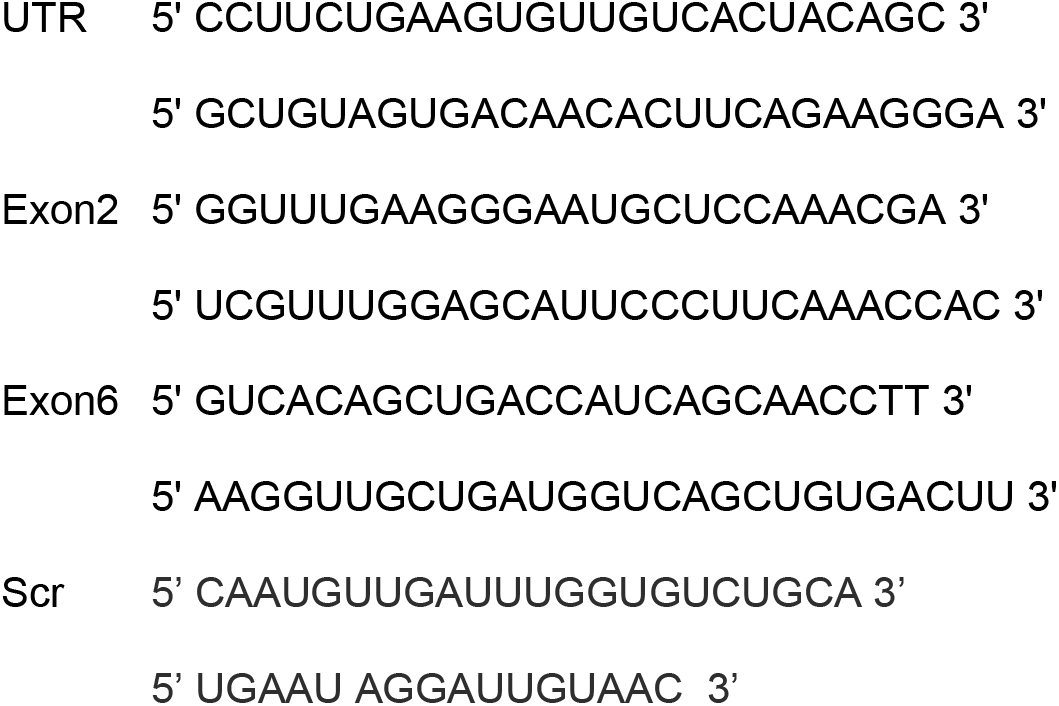

### Cell proliferation assay

Cell proliferation was performed as previously described^29^. Cells were plated in a 24-well plate at two densities of 5,000 and 10,000 cells per well in duplicate and cultured overnight. The following morning, plates were transferred to an incubator equipped with an IncuCyte^®^ S3 Live-Cell Analysis system (Essen BioSciences Inc, USA) for 6 days. Cell confluency was analyzed using the in-built IncuCyte^®^ S3 software.

### Clonogenic assays

Cells were plated on a 6-well plate at a density of 500 and 1000 cells per well and incubated for 14 days to determine colony viability. Colonies were fixed with 0.05% crystal violet for 30 minutes, washed and quantified for crystal violet colony counting and measurement by imaging on a GE InCell 2000 microscope and analysis by GE InCell 2000 3-D Deconvolution Software (GE Health care, Life Sciences, USA) and total surface area quantified by ImageJ v1.53q.

### Transwell assay

To measure cell migration 10^4^ cells were seeded 0.05% FBS onto the top of a transwell insert with 8 μM pores (Corning Inc. New York, USA) placed into a 24-well cell culture dish where 20% FBS was used as a chemoattractant in the base of the culture dish.Cells were incubated in the transwell plate at 37°C for 20 h and the migrated cells on the lower surface fixed at −20°C in ice cold MeOH for 30 min. After fixing cells were stained with Crystal violet (0.5% (w/v) in 25% MeOH (v/v)) for 30 min. To quantitate cell migration 6 fields of view per membrane were photographed with EVOS™ FL Auto 2 Imaging System(Thermo Scientific™ Invitrogen™), followed by quantification by ImageJ v1.53q (National Institutes of Health, USA).

### Immunoblotting

Cells were prepared for lysis as described previously^28^ with indicated antibodies (table below). The Super Signal chemiluminescent ECL-plus (Amersham) was used for signal detection.

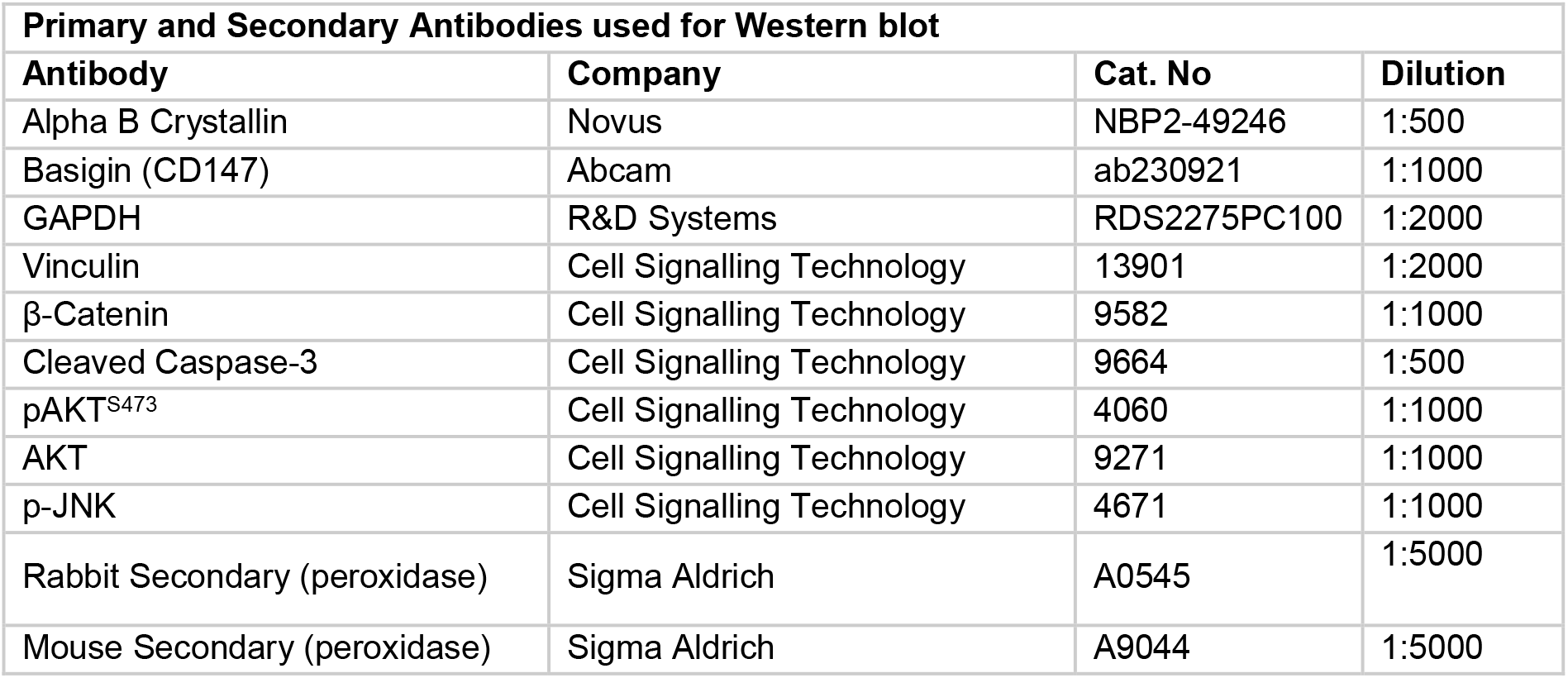

| Primary and Secondary Antibodies used for Western blot |  |  |  |
| --- | --- | --- | --- |
| Antibody | Company | Cat. No | Dilution |
| Alpha B Crystallin | Novus | NBP2-49246 | 1:500 |
| Basigin (CD147) | Abcam | ab230921 | 1:1000 |
| GAPDH | R&D Systems | RDS2275PC100 | 1:2000 |
| Vinculin | Cell Signalling Technology | 13901 | 1:2000 |
| β-Catenin | Cell Signalling Technology | 9582 | 1:1000 |
| Cleaved Caspase-3 | Cell Signalling Technology | 9664 | 1:500 |
| pAKT<sup>S473</sup> | Cell Signalling Technology | 4060 | 1:1000 |
| AKT | Cell Signalling Technology | 9271 | 1:1000 |
| p-JNK | Cell Signalling Technology | 4671 | 1:1000 |
| Rabbit Secondary (peroxidase) | Sigma Aldrich | A0545 | 1:5000 |
| Mouse Secondary (peroxidase) | Sigma Aldrich | A9044 | 1:5000 |

### Polymerase chain reaction

PCR was performed as described previously^28^ using Genomic DNA from animal tissue or cell pellet was extracted using QuickExtract DNA solution (Gene target solutions, QE09050).

### Immunofluorescence (IF)

IF was performed as described previously^28^ Briefly, 50,000 cells were seeded on UV pre-treated coverslips within a 24-well plate. After heat shock (43 C), to induce Cryab expression, cells were fixed using 4% Paraformaldehyde (PFA) followed by permeabilization (0.5% Triton X-100 for 5 min) and blocking (2% BSA in PBS) before staining with primary and secondary antibodies(Table 2 below) and mounting on the glass cover slides using Prolong Gold antifade reagent (Thermo Fisher Scientific). All immunofluorescence experiments were imaged using a DeltaVision Deconvolution Microscope (Applied Precision).

| Primary and Secondary Antibodies used for IF |  |  |  |
| --- | --- | --- | --- |
| Antibody | Company | Cat. No | Dilution |
| Alpha B Crystallin | Novus | NBP2-49246 | 1:200 |
| Basigin (CD147) | Abcam | ab230921 | 1:200 |
| Anti-mouse conjugated Alexa Fluor 488 | Thermofisher | A21202 | 1:600 |
| Anti-rabbit conjugated Alexa fluor 647 | Thermofisher | A31573 | 1:600 |
| DAPI | Sigma | D9542-5MG | 1:1000 |

### Mass-spectrometry

CryabWt and CryabTg MEFs were treated for 1h at 43°C to generate heat-shock stress before processing samples in triplicates for in-situ protein digestion as per previously described method^30^. For mass-spectrometry samples were loaded on to a Waters M-Class SYM100 trap column (180 um x 20 mm ID) for 6 min at a flow rate of 5 ul/min with 95% Solvent A (0.1% FA in water), and subsequently separated on a Waters BEH130 analytical column (75 um x 200 mm ID). Columns were equipped on a Waters nanoACQUITY UPLC coupled with a Thermo Orbitrap Fusion mass spectrometer. The solvent gradient ran at 300 nl/min and started at 92% Solvent A before ramping up to 27% Solvent B (0.1% FA in acetonitrile) over 45 min. This was followed by column washing and reequilibration for a total run time of 60 min. MS spectra were acquired in the mass range =350-1800 m/z (orbitrap resolution =60,000). Fragmentation for MS/MS spectra were acquired in the orbitrap at a resolution of 15,000 with a collision energy 30. The AGC target was 5e4, with a maximum ion injection time of 40 ms. The isolation window was set to 1.2 m/z. Dynamic exclusion was set to 15 sec and precursors with charge states from 2-6 were accepted for fragmentation.

Raw LCMS data was searched for protein IDs against the reviewed Uniprot mouse database (21,963 sequences, downloaded 24/04/2020) using Sequest HT on the Thermo Proteome Discoverer software (Version 2.2). Precursor and fragment mass tolerance were set to 20 ppm and 0.05 Da respectively. A maximum of two missed cleavages were allowed. A strict false discovery rate (FDR) of 1% was used to filter peptide spectrum matches (PSMs) and was calculated using a decoy search Carbamidomethylation of cysteines was set as a fixed modification, while oxidation of methionine and deamidation of glutamine and asparagine were set as dynamic modifications. Protein abundance was based on intensity of the parent ions and data was normalized based on total peptide amount.

Differentially expressed proteins were ranked based on upregulated ones over log_2_(0.6) or downregulated under log_2_(−0.6) in *Cryab*^Tg^ MEFs against *Cryab*^Wt^ MEFs. The biological pathway enrichment analysis was performed using GO analysis, and MCL cluster annotations and visualized using Cytoscape^31^ v.3.8.1

### CRYAB Expression Analyses

In human tumors, *CRYAB* gene expression and gene expression signature analyses were performed in samples from The Cancer Genome Atlas (TCGA) as described in the supplementary methods. All other CRYAB expression analyses were performed using samples from various datasets from the Gene Expression Omnibus (GEO, https://www.ncbi.nlm.nih.gov/geo) with the following accession numbers: normal cervix and early-stage cervical cancers (GSE7410), primary and metastatic renal cell carcinoma (GSE31232), primary tumours without and with metastasis in hepatocellular carcinoma (GSE45114), pancreatic cancer (GSE63124), breast cancer (GSE9893) and colorectal cancer (GSE87211), locally and distantly metastasized pancreatic cancer (GSE34153), prostate cancer (GSE74367) and lung cancer (GSE18549).

### DMBA tumorigenic treatments

DMBA (7,12-Dimethylbenz[a]anthracene) treatments consisted of a single administration of 50 μl of a solution 0.5% DMBA (Sigma) to the dorsal surface on postnatal day 5 of mice.

### Statistical Analyses

All statistical analyses were performed using GraphPad Prism v 8.0, using a general linear statistical model, as defined in each section. The error bar represents the mean ± Standard Deviation (SD) unless indicated otherwise. The statistical significance of the *p*-value is designated with an asterisk (*); *p*-values: * *p* < 0.05, ** *p* < 0.01, *** *p* < 0.001, and **** *p* < 0.0001.

## Acknowledgments

We acknowledge all members of the Signal transduction laboratory for their critical comments and advice, Debottam Sinha for his contribution for analysis of control cohort as well as Rebekah Ziegman in processing proteomics data and Alejandro Lopez for editing and commenting on some parts of this manuscript.

## Author Contributions

Conceptualization: JS, ALB, BR, KKK; Methodology: JS, ALB, BR, KKK; Investigation: ALB, BR, SMT, CS; Bioinformatics: PD, PMA, SMT; Pathology: JF, SS; Writing the original draft preparation: SMT, BR, KKK; Writing review and editing: BR, ALB, PMA, KKK; supervision: ALB, BR, KKK. All authors have read and agreed to the published version of the manuscript.

---

**Supplementary Table 1**: Correlation analysis.

**Supplementary Table 2**: List of the upregulated genes and annotation.

## Supplementary Figures

**Fig S1.**
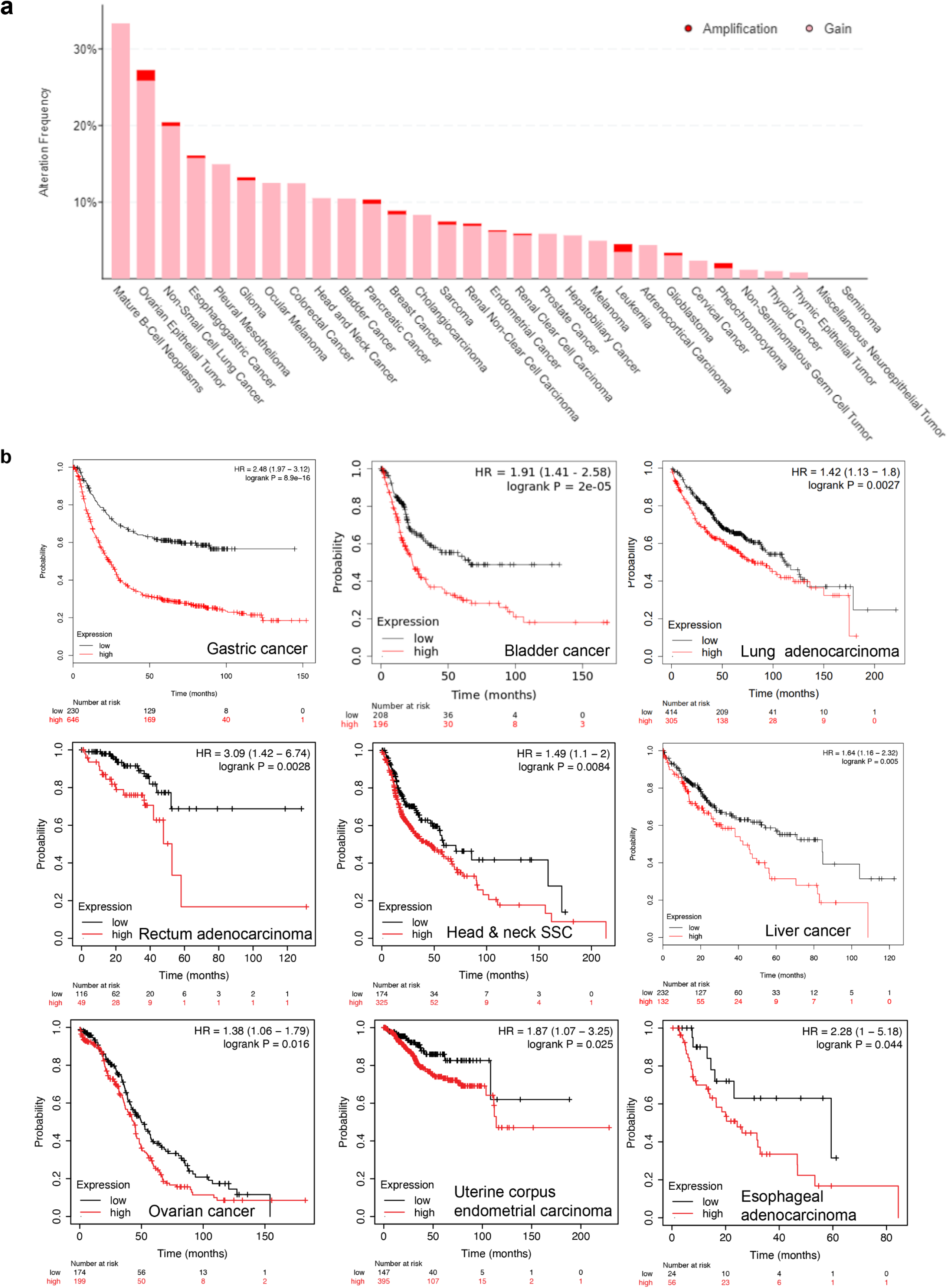
*CRYAB* is frequently gained and high expression is associated with poor survival in multiple cancer types. **(a)** Chart shows the percentages of copy number amplification (red) or copy number gain (pink) of *CRYAB* across various cancers using the cBioPortal for Cancer Genomics. **(b)** Kaplan-Meier survival analysis for TCGA cancer type indicated below the chart for the overall survival. Patients were split into two groups of high (red) and low (black) *CRYAB* expression levels. Numbers of patients at risk are shown at the bottom.

**Fig S2.**
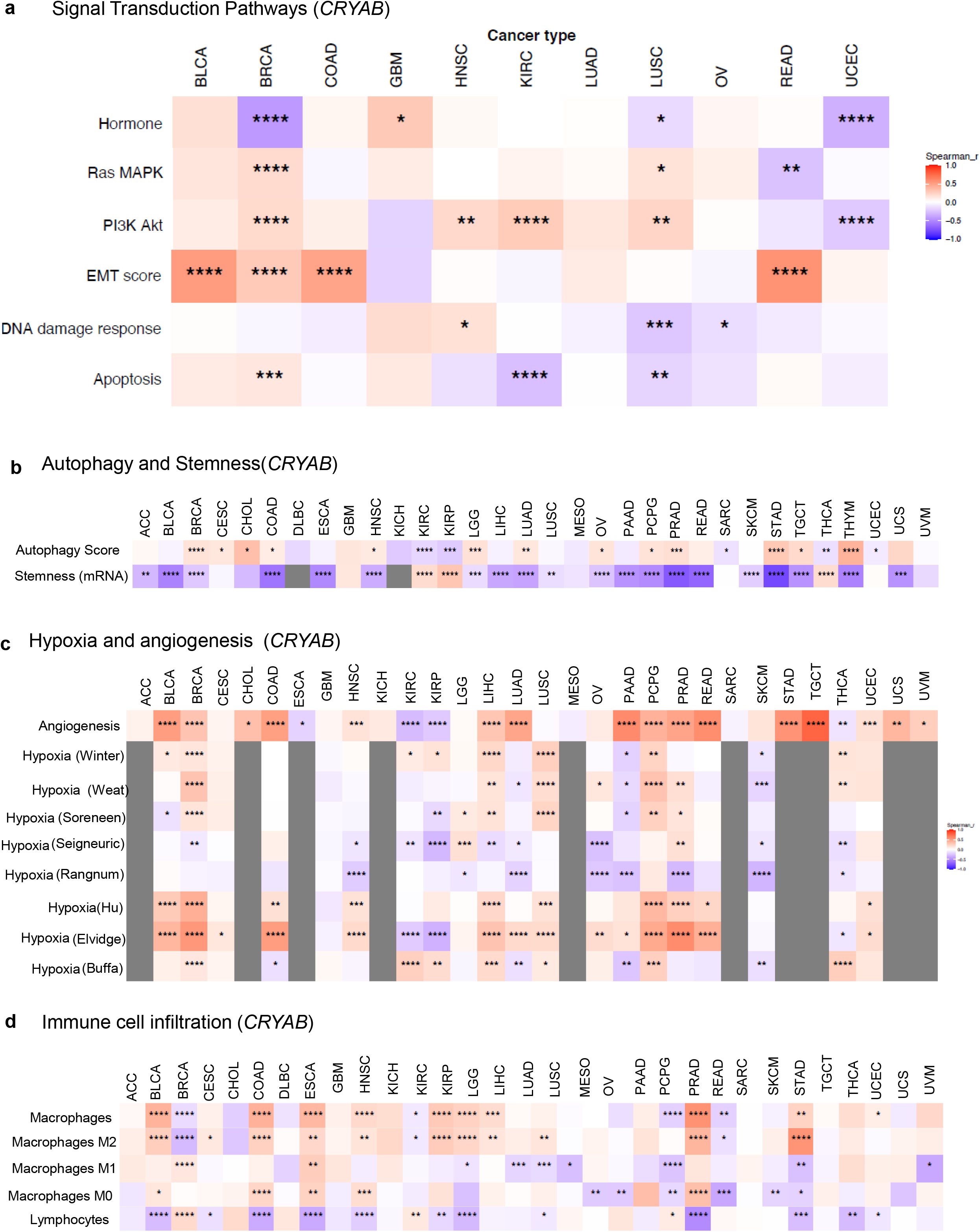
*CRYAB* correlated with deregulation of signaling, oncogenic features and tumor infiltration across some cancers. **(a-d)** The heatmaps show the correlations between CRYAB gene expression levels and variables (determined via the integration of multi-platform data from several types of ‘omics’ analyses) listed on the y-axis in up to 32 cancer types, whose abbreviations are listed on the top x-axis based on the methodology previously described^29^; for all analysis **(a-d)**: each tile in the heatmaps shows the Spearman correlation between the gene expression levels (mRNA levels from The Cancer Genome Atlas (TCGA) RNAseq datasets) and the variables listed below. The color of each tile reflects the Spearman correlation coefficient (r), as indicated by the color key on the right. Each tile shows the Spearman p value abbreviations in the following format: blank tiles: p > 0.05; *, p < 0.05; **, p < 0.01; ***, p < 0.001; ****, p < 0.0001. <u>Cancer type abbreviations</u>: ACC, Adrenocortical Cancer; BLCA, Bladder Cancer; BRCA, Breast Cancer; CESC, Cervical Cancer; CHOL, Cholangiocarcinoma (bile duct cancer); COAD, Colon Cancer; DLBC, Large B-cell Lymphoma; ESCA, Esophageal Cancer; GBM, Glioblastoma; HNSC, Head and Neck Cancer; KICH, Kidney Chromophobe; KIRC, Kidney Clear Cell Carcinoma; KIRP, Kidney Papillary Cell Carcinoma; LGG, Lower Grade Glioma; LIHC, Liver Cancer; LUAD, Lung Adenocarcinoma; LUSC, Lung Squamous Cell Carcinoma; MESO, Mesothelioma; OV, Ovarian Cancer; PAAD, Pancreatic Cancer; PCPG, Pheochromocytoma & Paraganglioma; PRAD, Prostate Cancer; READ, Rectal Cancer; SARC, Sarcoma; SKCM, Melanoma; STAD, Stomach Cancer; TGCT, Testicular Cancer; THCA, Thyroid Cancer; THYM, Thymoma; UCEC, Endometrioid Cancer; UCS, Uterine Carcinosarcoma; UVM, Ocular melanoma.

**Fig S3.**
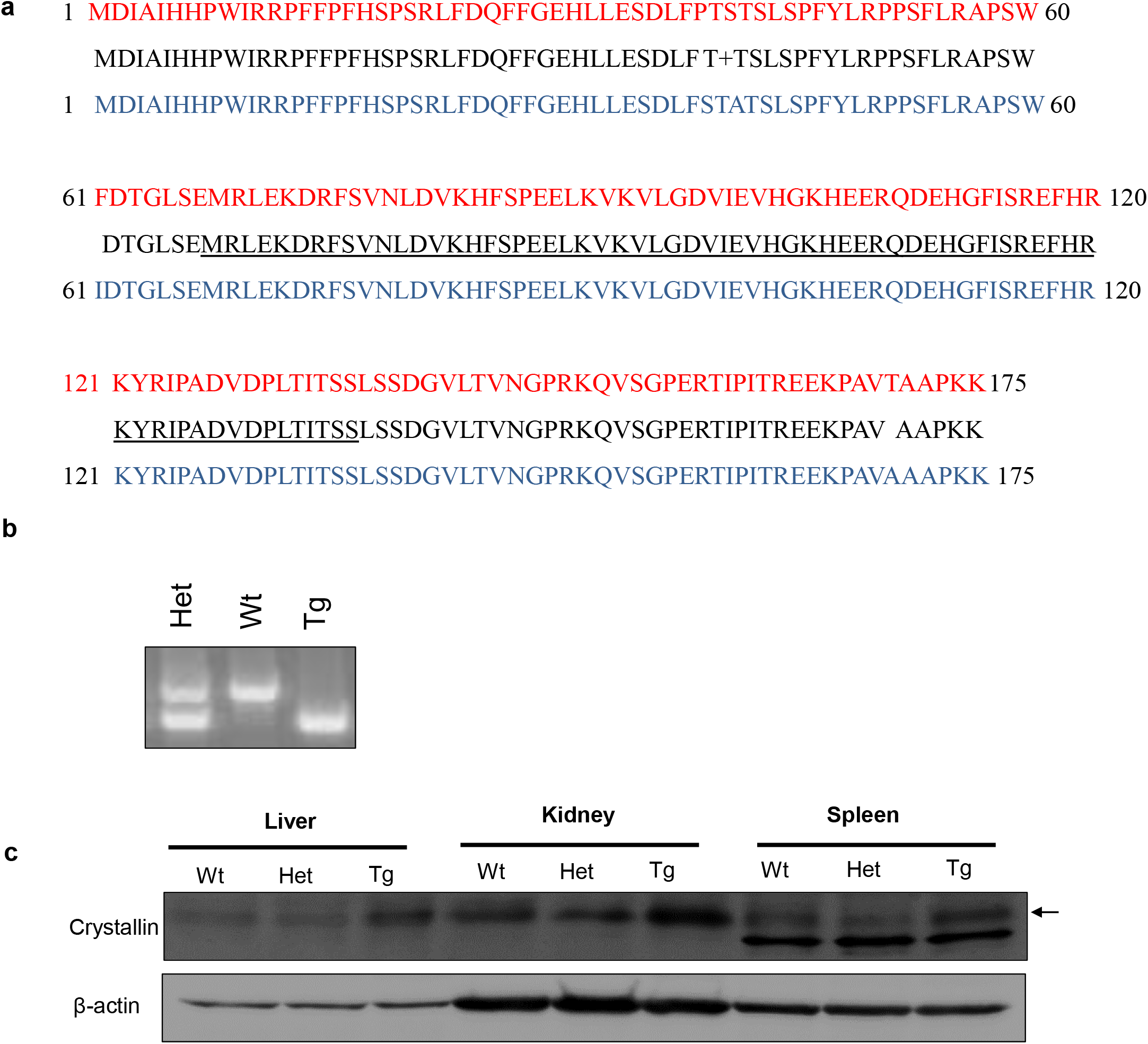
Protein alignment and mouse model target validation. **(a)** Alignment of human (NP_001876, upper in red) and mouse (NP_034094.1, lower in blue) αB-Crystallin protein sequences. The spaces indicate sequence mismatch and a + indicates a positive match on the scoring matrix and αB-crystallin domain is underlined in the consensus. **(b)** PCR genotyping showing *Cryab*^Wt/Tg^, *Cryab*^Wt/Wt^ and *Cryab*^Tg/Tg^ genotypes. **(c)** Immunoblot analysis of Crystallin protein expression of *Cryab*^Wt/Wt^, *Cryab*^Wt/Tg^, and *Cryab*^Tg/Tg^ mouse tissues as indicated in the figure. β-actin was used as a loading control.

**Fig S4.**
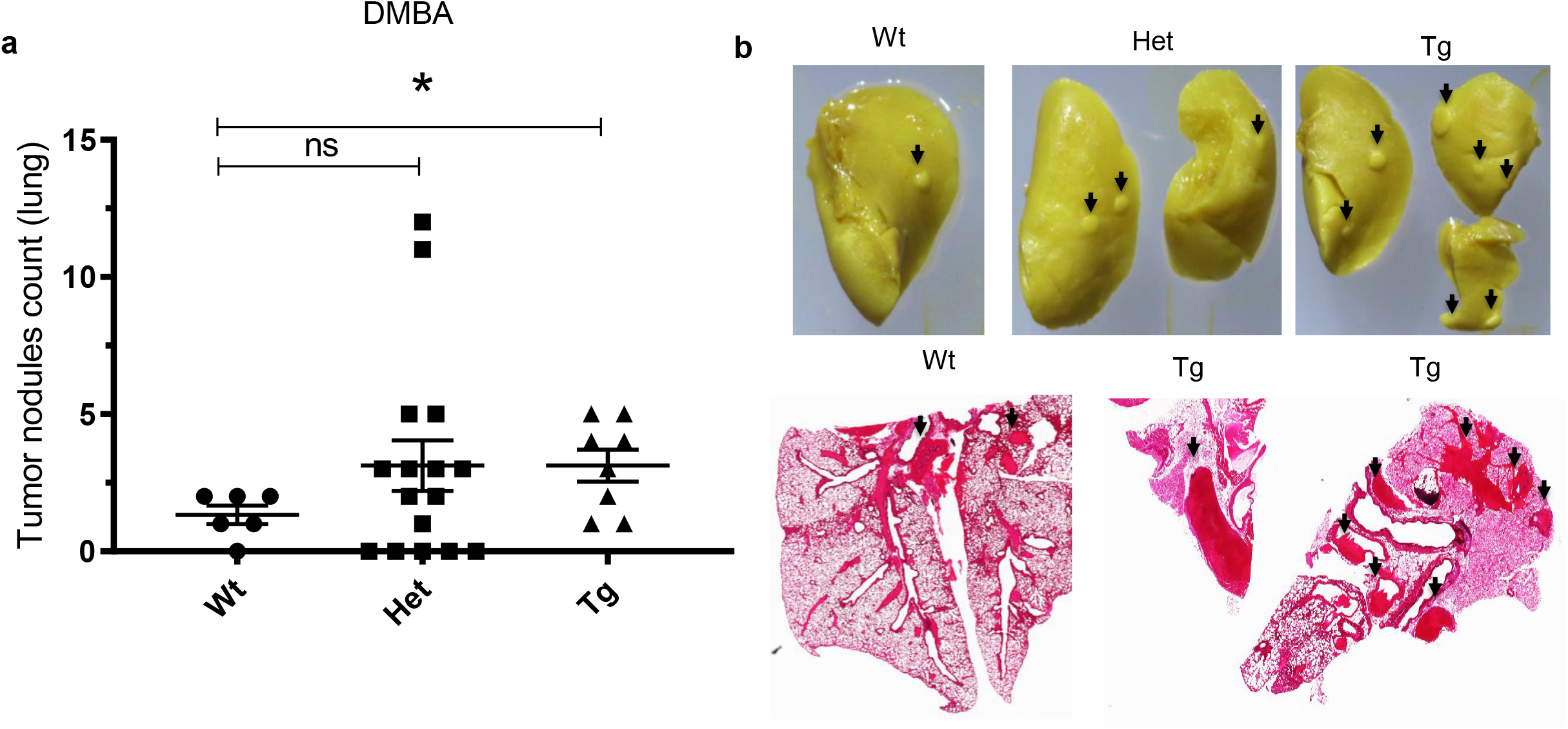
DMBA experiment. **(a)** Chart shows the counted number of nodules from dissected lungs of DMBA treated *Cryab*^Wt/Wt^, *Cryab*^Wt/Tg^, and *Cryab*^Tg/Tg^ the outliers were removed using the ROUT method (Q = 1%) based on the False Discovery Rate (FDR). **(b)** representative image of Bouin’s solution fixed lungs from each genotype (upper) and H&E stained tissue from *Cryab*^Wt/Wt^ and Cryab^Tg/Tg^ (lower), arrows show the nodules or tumors, notice to the bigger size of tumors in *Cryab*^Tg^

**Fig S5.**
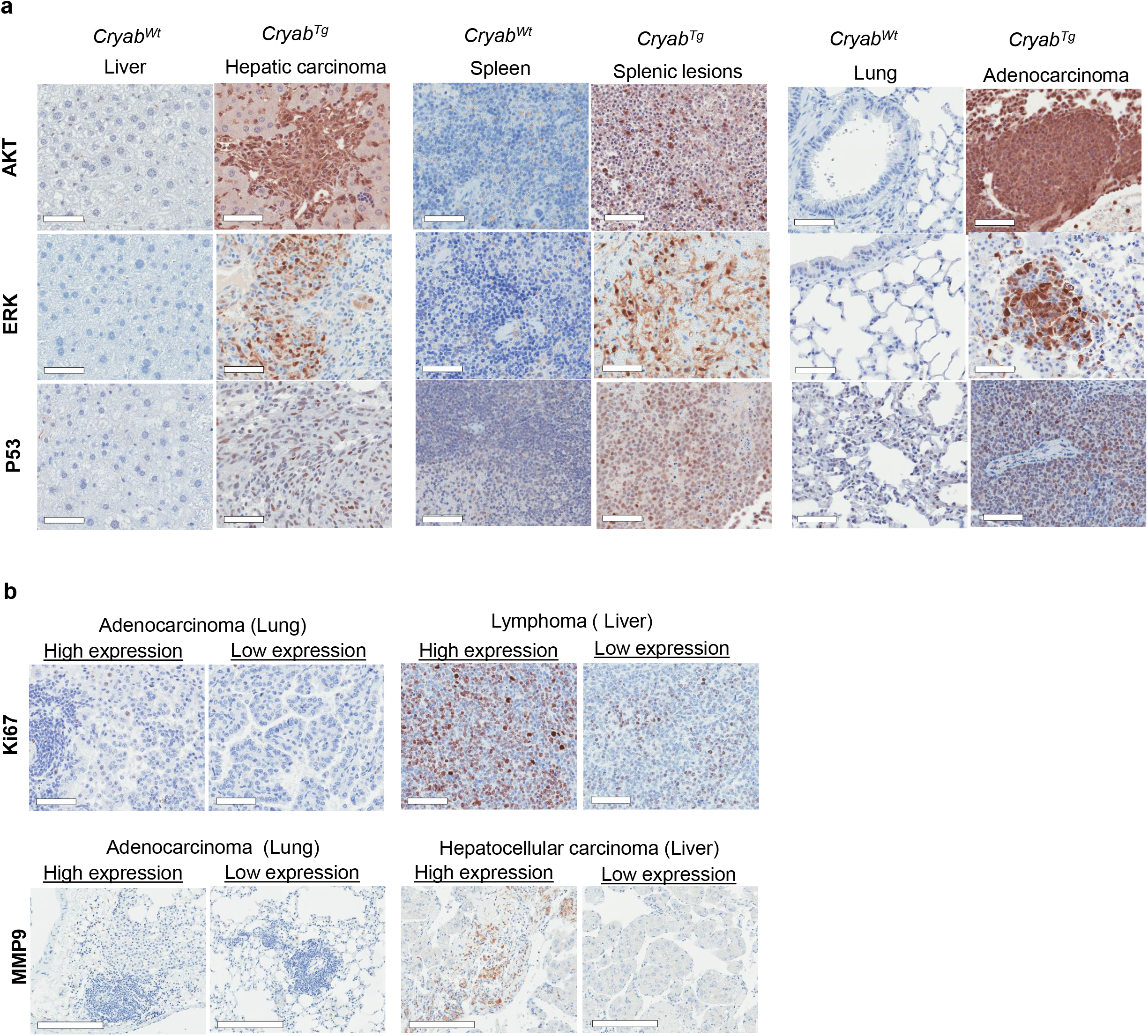
IHC on tumors and normal tissues. **(a)** IHC staining compares the expression level of indicated markers in malignant tissue of *Cryab*^Tg/Tg^ with age and tissue matched control from *Cryab*^Wt/Wt^, **(b)** IHC staining compares the high and low expression level of indicated markers in malignant tissue of *Cryab*^Tg^ as shown in the figure.

**Fig S6.**
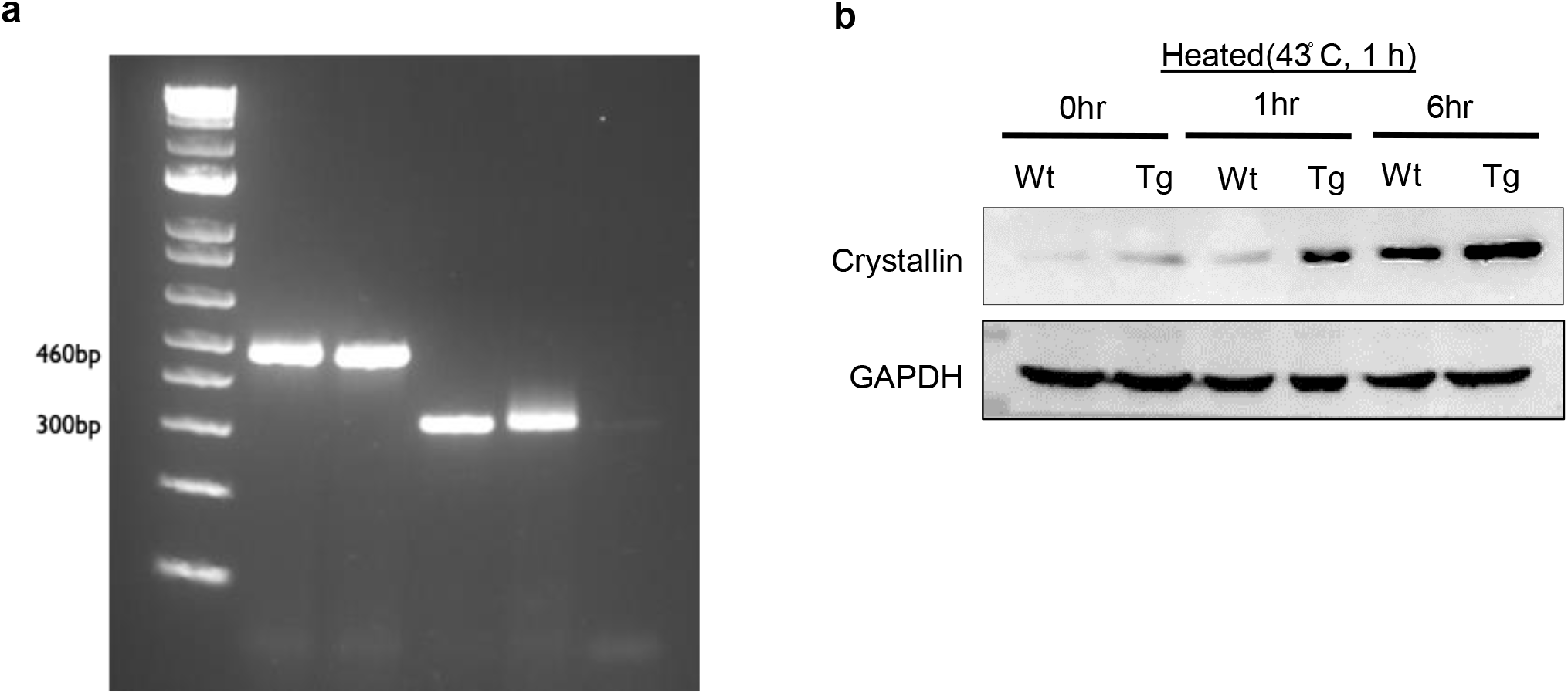
Genotyping and WB on MEFs. **(a)** PCR genotyping showing *Cryab*^Wt/Wt^ (Wt) and *Cryab*^Tg/Tg^ (Tg) E1A/Ras transformed MEFs genotypes **(b)** Immunoblot analysis of Crystallin protein expression of *Cryab*^Wt/Wt^, and *Cryab*^Tg/Tg^ E1A/Ras transformed MEFs after heat-shock (43° C for 1 hour) following by time course incubation as indicated in the figure. GAPDH was used as a loading control.

**Fig S7.**
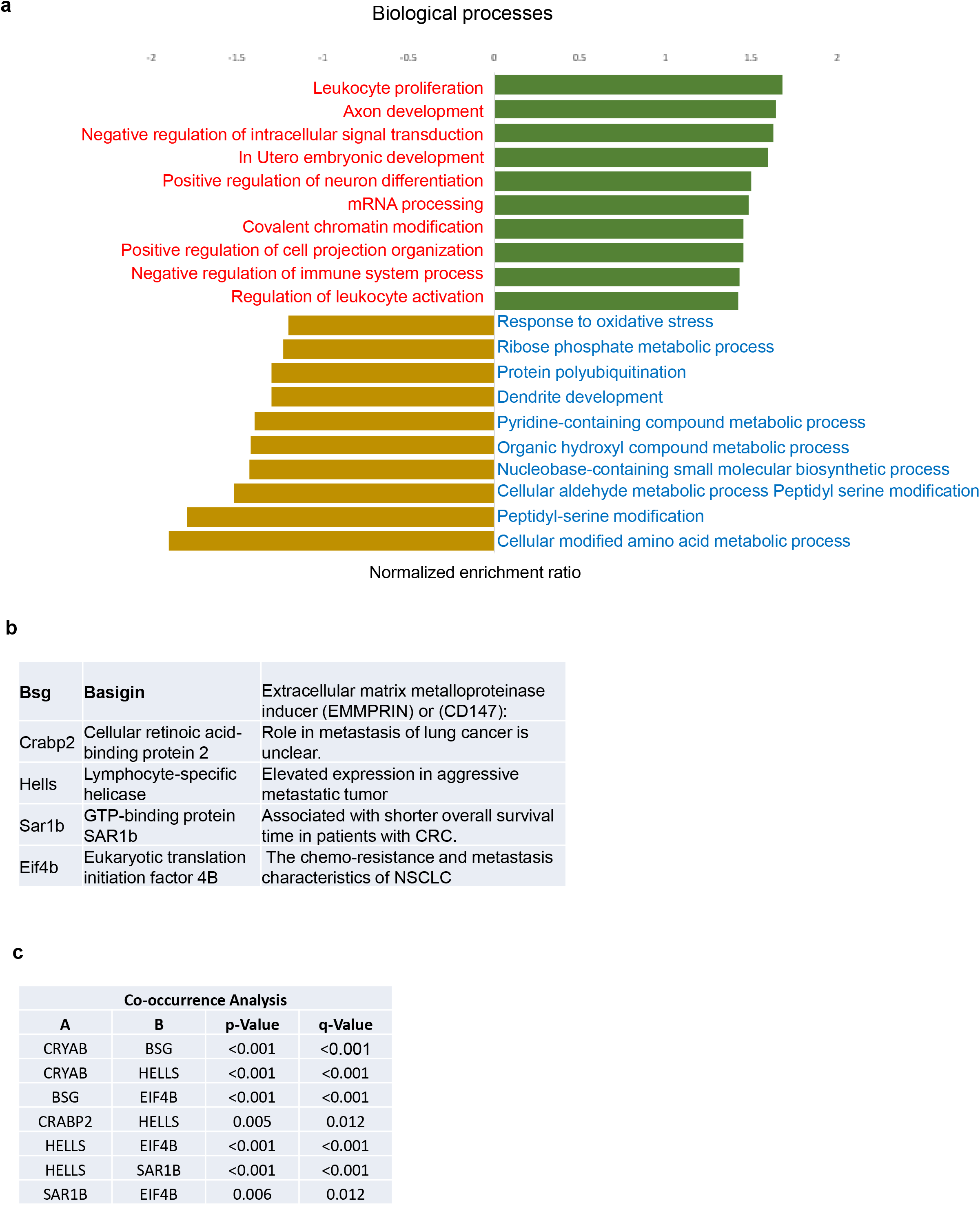
Biological process annotation. **(a)** Functional annotation and pathway enrichment analysis of differentially regulated proteins of the proteomic data comparing Cryab Tg E1ARas-transformed MEFs to Cryab WT MEFs for either upregulated (green) or downregulated (yellow), Data were normalized to the enrichment ratio. Data was plotted using default parameters of DAVID (the database for annotation, visualization and integrated discovery) **(b)** The table shows the top five highest upregulated genes. **(c)** co-occurance analysis of CRYAB gene and the top five highest upregulated genes from TCGA pan-cancer

**Fig S8.**
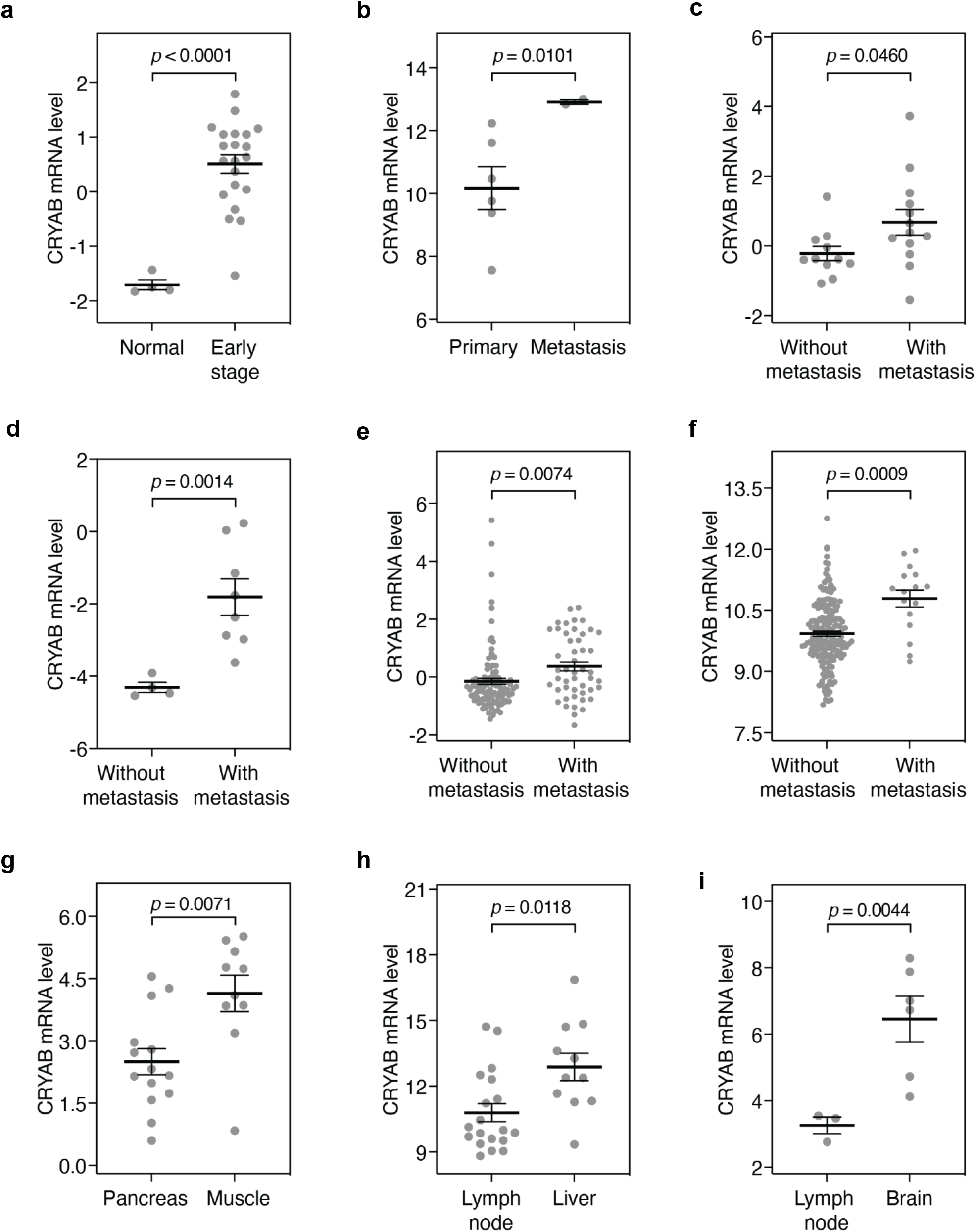
CRYAB expression is associated with metastasis in human tumors. **(a)** CRYAB expression levels in normal cervix and early-stage cervical tumors. **(b)** CRYAB expression levels in primary and metastatic renal cell carcinoma. **(c-f)** CRYAB expression levels in primary tumors without and with metastasis in, respectively: **(c)** hepatocellular carcinoma, **(d)** pancreatic cancer, **(e)** breast cancer and **(f)** colorectal cancer. **(g)** CRYAB expression levels in pancreatic cancer locally metastasized to pancreas or distantly metastasized to muscle. **(h)** CRYAB expression levels in prostate cancer locally metastasized to lymph node or distantly metastasized to liver. **(i)** CRYAB expression levels in lung cancer locally metastasized to lymph node or distantly metastasized to brain. Error bars indicate means and standard errors. *P* values: two-sided *t* tests.

## Supplementary methodology

### TCGA data analysis

#### Signal Transduction Pathways

Pathway scores for each of these eight pathways were determined by integration of multi-platform data from several types of ‘omics’ analyses, as described by Hoadley et al., 2010 (PMID 25109877).

Hypoxia and angiogenesis: Angiogenesis score: A 43-gene expression signature for the level of angiogenesis, as determined by Masiero et al., 2013 (PMID 23871637). Hypoxia score (Winter): A 99-gene expression signature for the level of hypoxia, as determined by Winter et al., 2007 (PMID 17409455). Hypoxia score (West): A 26-gene expression signature for the level of hypoxia, as determined by Eustace et al., 2013 (PMID 23820108). Hypoxia score (Sorensen): A 27-gene expression signature for the level of hypoxia, as determined by Sorensen et al., 2010 (PMID 20429727). Hypoxia score (Seigneuric): A gene expression signature for the level of hypoxia, as determined by Seigneuric et al., 2007 (PMID 17532074). Hypoxia score (Ragnum): A 32-gene expression signature for the level of hypoxia, as determined by Ragnum et al., 2015 (PMID 25461803). Hypoxia score (Hu): A 13-gene expression signature for the level of hypoxia, as determined by Hu et al., 2009 (PMID 19291283).

Autophagy and stemness: Autophagy-related prognostic signature: A prognostic gene expression signature based in the weighted expression of 8 autophagy-related genes, as described by An et al., 2018 (PMID 30410611). Stemness (mRNA): Level of stemness, based on mRNA data, determined as described by Malta et al., 2018 (PMID 29625051).

Immune cell infiltration: The levels of indicated tumour-infiltrated immune cell types were estimated using the tumor immune estimation resource, developed by Li et al., 2016 (PMID 27549193).

