## Supplementary material for "Alpha-B-Crystallin overexpression is sufficient to promote tumorigenesis and metastasis in mice": Sup: Table 1.docx

| Malignancies | Incidence  (frequency) | Features |
| --- | --- | --- |
| Hemangiosarcoma | (30%) | This neoplasm was comprised of vascular spaces of varying size, sometimes containing thrombi, and lined by a pleomorphic population of proliferating endothelial cells with frequent mitotic figures present. |
| Hepatic Carcinoma (hepatocellular carcinoma) | (25%) | There was trabecular growth in irregularly thick plates of neoplastic hepatocytes, which sometimes resembled normal hepatocytes, but often had enlarged, hyperchromatic nuclei, prominent nucleoli, and abundant cytoplasm (megalocytosis). |
| Alveolar/Bronchiolar carcinoma | (10%) | In the lung, there was papilliform growth of a pleomorphic population of epithelial cells, which were arranged around a fibrovascular core. |
| B-cell lymphoma | (10%) | There was effacement of the normal splenic architecture by a proliferating population of CD79a-immunopositive lymphoblastic cells. |
| Histiocytic sarcoma | (5%) | Tumour was composed of sheets of large, pleomorphic histiocytic cells with abundant cytoplasm; multinucleated giant cells were common. |
| Metastasis | (20%) | Spleen to liver and lung metastasis, liver (HCC) to lung and liver metastasis |

The distribution of cancer spectrum in *Cryab* transgenic mice.
